# Probing the basis of disease heterogeneity in multiple sclerosis using genetically diverse mice

**DOI:** 10.1101/2024.06.03.597205

**Authors:** Emily A. Holt, Anna Tyler, Taylor Lakusta-Wong, Karolyn G. Lahue, Katherine C. Hankes, Cory Teuscher, Rachel M. Lynch, Martin T. Ferris, J. Matthew Mahoney, Dimitry N. Krementsov

## Abstract

Multiple sclerosis (MS) is a complex disease with significant heterogeneity in disease course and progression. Genetic studies have identified numerous loci associated with MS risk, but the genetic basis of disease progression remains elusive. To address this, we leveraged the Collaborative Cross (CC), a genetically diverse mouse strain panel, and experimental autoimmune encephalomyelitis (EAE). The thirty-two CC strains studied captured a wide spectrum of EAE severity, trajectory, and presentation, including severe-progressive, monophasic, relapsing remitting, and axial rotary (AR)-EAE, accompanied by distinct immunopathology. Sex differences in EAE severity were observed in six strains. Quantitative trait locus analysis revealed distinct genetic linkage patterns for different EAE phenotypes, including EAE severity and incidence of AR-EAE. Machine learning-based approaches prioritized candidate genes for loci underlying EAE severity (*Abcc4* and *Gpc6*) and AR-EAE (*Yap1* and *Dync2h1*). This work expands the EAE phenotypic repertoire and identifies novel loci controlling unique EAE phenotypes, supporting the hypothesis that heterogeneity in MS disease course is driven by genetic variation.

**Summary:** The genetic basis of disease heterogeneity in multiple sclerosis (MS) remains elusive. We leveraged the Collaborative Cross to expand the phenotypic repertoire of the experimental autoimmune encephalomyelitis (EAE) model of MS and identify loci controlling EAE severity, trajectory, and presentation.

## Introduction

Multiple sclerosis (MS) is an autoimmune disease of the central nervous system (CNS) characterized by demyelination, gliosis, axonal loss, and progressive neurological dysfunction, representing the leading cause of non-traumatic neurological disability in young adults^1^. The pathogenesis of MS is not fully understood, but current evidence suggests that activation of myelin-reactive CD4 T cells triggers an inflammatory cascade in the CNS, recruiting other immune cells, which mediate subsequent tissue destruction and pathology^2,3^. Significant heterogeneity in disease presentation and severity exists, with disease courses characterized as: (i) clinically isolated syndrome (CIS), denoted as the first inflammatory and demyelinating event preceding MS diagnosis^4^; (ii) relapsing remitting MS (RR-MS), defined by distinct flares of disease progression with minimal disease progression between relapses and an overall less severe disease course; (iii) secondary progressive MS (SP-MS), which typically transitions into RR-MS and presents as a progressive deterioration over time; and (iiii) primary progressive MS (PP-MS), which presents as a severely debilitating disease course with early disease progression and poor prognosis^5^. In addition, significant sex differences have also been noted, with MS incidence being approximately three times higher in women, while disease course has been shown to be more severe in men^6,7^. The biological underpinnings behind this heterogeneity in disease presentation remain largely unknown, and their characterization should yield prognostic indicators to better inform personalized therapeutic intervention^8^.

Previous studies have shown that up to 30% of MS risk can be attributed to genetic factors^9^. Early studies in MS families demonstrated a significant genetic component, which was mapped to the major histocompatibility (MHC) locus, with HLA-DRB1*15:01 being the strongest risk allele^10,11^. Nonetheless, the MHC locus accounted for only a portion of the total heritable risk for this polygenic disease. Subsequent genome-wide association studies (GWAS) have identified 200 non-MHC loci, as well as 32 independent loci within the MHC, associated with MS incidence^9^. However, despite the strong genetic association with MS disease risk, the genetic basis for MS disease course heterogeneity remains poorly understood. An additional limitation of these association studies lies in that the causality of the identified genes cannot be demonstrated in humans. In this regard, mouse models have been proposed as the next crucial step in the post-GWAS era^12^.

Several mouse models of MS exist, with experimental autoimmune encephalomyelitis (EAE) being the principal immune-mediated model of this disease. This model has been instrumental in improving our understanding of MS pathogenesis at the cellular and molecular levels, as well as developing new disease modifying therapies^13,14^. However, animal models of MS, in particular EAE, have been criticized for failing to capture many relevant aspects of the human disease^15,16^. We propose that this shortcoming is in part due to the failure to consider the importance of genetic heterogeneity that is so pronounced in human populations; a gap that we attempted to address in this study. With regard to genetics approaches in animal models, the most common approach has been to use reverse genetics to delete a candidate gene of interest on a fixed genetic background, typically C57BL/6 (B6). While effective, such genetically modified mice are rarely representative of natural genetic variation present in the human population^17^, where variation in many native genomic elements regulates the expression and activity of genes in a time- and tissue-specific manner. Additionally, the candidate gene is pre-selected based on prior knowledge. In contrast, unbiased forward genetics approaches utilize natural genetic variants that control a phenotype of interest, analogous to allelic variants naturally segregating in human populations^18^. Genetic crosses between two or more populations of interest and subsequent mapping can reveal loci containing genes linked to the phenotype, akin to GWAS in humans. Importantly, the causal effects of candidate genes can be confirmed using a number of approaches, including congenic mapping/positional cloning, targeted genome editing, transgenic complementation, and pharmacological targeting^12^.

While conventional laboratory inbred strains of mice are an important tool in forward genetics, they represent artificially selected organisms originating from a small founder population^19^. As such, they lack the range of genetic diversity and selective evolutionary pressure present in human populations. These limitations can be overcome by incorporating into the experimental design so-called wild-derived inbred strains^20^, thereby more accurately modeling human genetic diversity. This approach has been used previously in our lab by leveraging wild-derived PWD/PhJ (PWD) mice^21^. Our initial studies showed that PWD mice have a decreased susceptibility to EAE, associated with a downregulation of proinflammatory and MS-associated genes in immune cells^22^. To begin to map loci driving these phenotypes, we employed the B6.Chr^PWD^ chromosome substitution (consomic) strain panel, which carry PWD chromosomes on the B6 background^23^. Using B6.Chr^PWD^ consomic mice, we demonstrated that PWD-derived alleles profoundly regulated EAE severity, often in a sex-specific manner^24,25^. While B6.Chr^PWD^ consomic mice could be used as a starting point for mapping of specific gene variants driving EAE phenotypes (e.g. using congenic mapping), this is a laborious process. Further, the genetic diversity, although improved, is still limited to 2 allelic variants per gene (PWD and B6) and captured only classic EAE symptomatology (ascending paralysis). To capture a broader spectrum of disease heterogeneity more representative of MS, we turned to the Collaborative Cross (CC) mouse genetic resource. The CC is a panel of multi-parental recombinant inbred lines designed specifically for the analysis of complex and polygenic phenotypes. They represent a unique resource to discover and dissect the exact contributions of genetic, environmental, and developmental components to the etiology of common complex human diseases^26^. These mice were generated using 8 founder strains. Five of these are conventional laboratory inbred strains: C57BL/6J (B6), A/J, 129S1/SvImJ (129S1), NOD/ShiLtJ (NOD), NZO/HlLtJ (NZO), and three are wild-derived strains: CAST/EiJ (CAST), PWK/PhJ (PWK), and WSB/EiJ (WSB). Of note, the PWK strain is a close relative of the PWD strain used in our previous studies, as it was developed in parallel from the same wild-derived population^21^. Collectively, these founder genomes cover ∼90% of the known genetic variation in mice. Using a combinatorial breeding design, through a long series of intercrosses between these different strains and their offspring, the CC strains were generated and fixed as homozygotes, allowing for unlimited numbers of repeat phenotypic measurements in genetically identical individuals^26,27^.

Since becoming commercially available, the CC mice have been applied to several research fields, including examining the genetic susceptibility to infectious diseases and control of immunologic phenotypes^28–36^, as well as allergic responses^37–40^, but to our knowledge this model has not yet been applied to study organ-specific autoimmunity. Here, we developed an EAE induction protocol for use in CC mice, using strains carrying *H2^b^* and *H2^g7^* MHC haplotypes. Using this approach, we characterized EAE phenotypes in 32 different CC strains, including: EAE incidence/susceptibility, EAE subtype (classic or atypical), and overall disease course presentation. This revealed a wide variation in EAE phenotypes and identified several strains with clinically relevant phenotypes, including but not limited to: EAE resistance, severe progressive disease, relapsing remitting disease, monophasic disease, and atypical axial rotary (AR)-EAE, as well as the presence of genotype-specific sex differences. Subsequent follow up analysis revealed distinct immunopathology associated with EAE phenotypes of interest. Furthermore, utilization of this approach allowed for quantitative trait loci (QTL) mapping and subsequent candidate gene prioritization, revealing distinct linkage patterns and identifying novel loci and genes controlling unique EAE phenotypes. Taken together, this characterization greatly expands the phenotypic repertoire of the EAE model, bringing the model one step closer to human disease relevance by addressing the role of genetic diversity in disease presentation. Importantly, together with emerging genome-wide studies in humans^41,42^, these findings strongly support the hypothesis that heterogeneity in MS disease course is driven by natural genetic variation.

## Results

### MOG_35-55_ induced EAE in H2^b^ and H2*^g7^* Collaborative Cross (CC) strains captures a broad range of clinically relevant disease phenotypes

EAE, like MS, is initiated by autoreactive CD4 T cells recognizing myelin antigens presented on major histocompatibility complex (MHC) class II molecules. These autoreactive CD4 T cells are typically elicited by immunization with myelin antigens together with adjuvants such as complete Freund’s adjuvant (CFA) and pertussis toxin (PTX), with the latter likely serving to disrupt the blood-brain barrier^43^. Several different immunogens are used for EAE induction. These include crude mouse spinal cord homogenate (mSCH), recombinant myelin proteins, or, most commonly, peptides derived from myelin proteins, including myelin oligodendrocyte glycoprotein 35-55 peptide (MOG_35-55_) classically used in C57BL/6J (B6) mice^8–10^. It has been well documented that different strains of mice have varying susceptibility to the EAE peptide immunogens based on the ability of their MHC allelic variants to bind to and present these immunogens^44–46^. This is potentially problematic to the design of a universal EAE induction protocol for CC strains, as the original 8 CC founder strains have contributed 7 different MHC haplotypes to the resulting CC strains (**Table S1**). Given this, we initially attempted to induce EAE utilizing mSCH, to capture all potential neuroantigens and therefore provide potential antigen binding across various MHC allelic variants. However, initial experiments in a subset of CC strains demonstrated that mSCH-based induction, although fairly effective in B6 mice, had variable and low penetrance of EAE in CC mice (**Fig. S1**).

As an alternative, we chose a peptide-based EAE induction approach utilizing the classic MOG_35-55_ peptide, based on the identification of compatible allelic variants at the MHC locus (called *H2* in the mouse). Previous studies have shown successful using MOG_35-55_-based EAE induction in three classic laboratory strains, which were also used as founders for the CC: B6, 129S1/SvImJ (129S1), and NOD/ShiLtJ (NOD)^44–46^. These strains carry either an *H2^b^* (B6 and 129S1) or a *H2^g7^* (NOD) MHC haplotype (**Table S1**), both which would be expected to be found in the CC strains. Using the available CC genotype data (see Materials and Methods), we determined that a total of 32 out of the 69 commercially available CC strains carry an *H2^b^* or *H2^g7^* haplotype at MHC class II loci (**Table 1**). These 32 CC strains are thus potentially compatible with the MOG_35-55_ based EAE induction approach and were selected for inclusion in our study (**Fig. 1A**).

**Figure 1.**
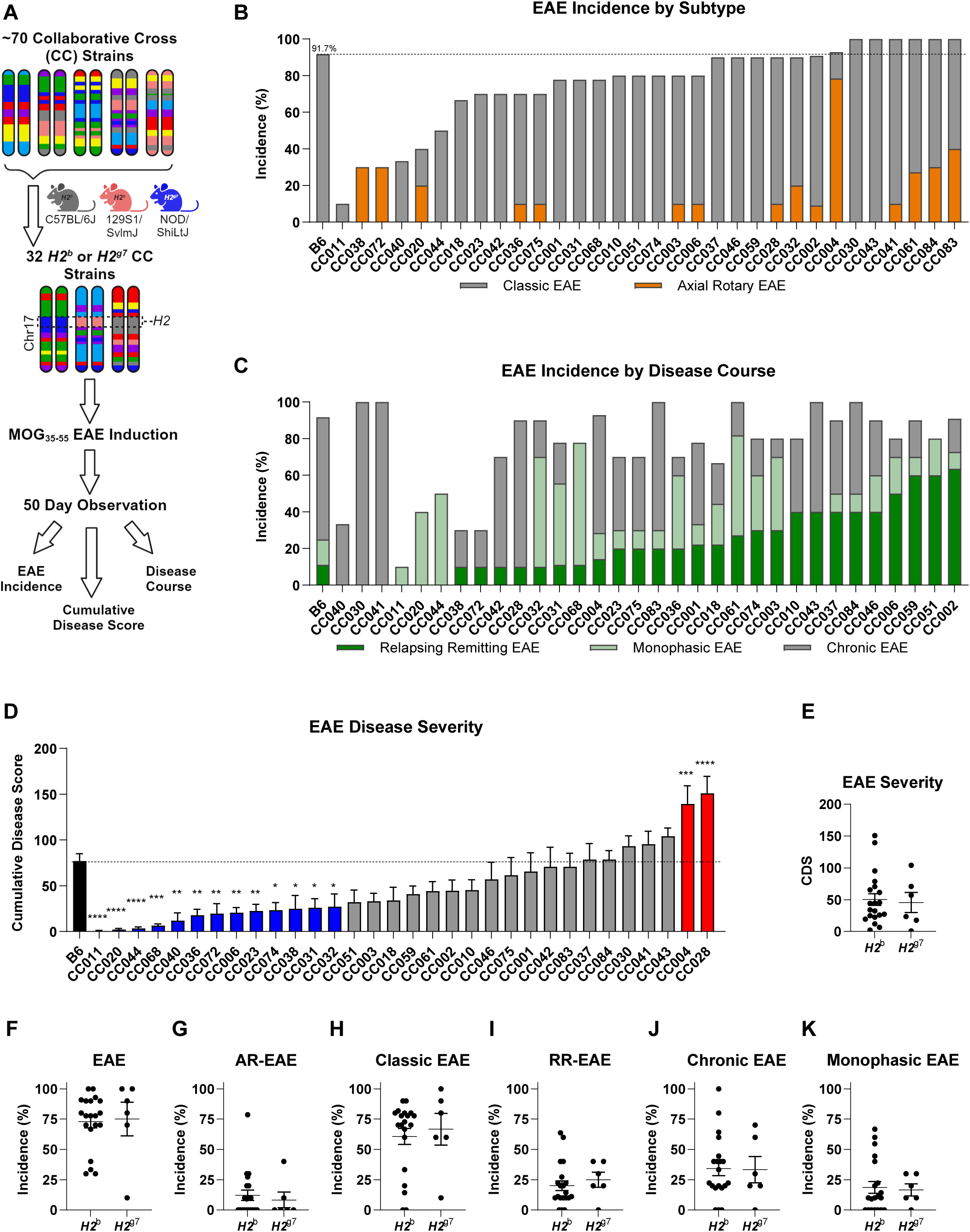
MOG_35-55_ induced EAE in CC strains results in heterogeneous disease profiles. EAE was induced in 8-13 week old male and female B6 and *H2^b^* or H2^g7^ CC mice (32 strains, ∼5 males and ∼5 females per strain) by s.c. immunization with 200 µg of MOG_35-55_ emulsified in CFA, and a single i.p. injection of 200 ng of PTX on D0 (see Materials and Methods). Mice were observed daily for a total of 50 days starting at 5 days post induction for the presence of clinical disease symptoms which were quantified to assess the overall EAE disease profile, as described in Materials and Methods. (**A**) Workflow illustrating the study design and selection of CC strains compatible with MOG_35-55_ EAE induction for inclusion based on founder MHC haplotype (*H2^b^* or *H2^g7^*). (**B**) Percent EAE incidence per strain, in CC strains, with B6 mice shown as a reference control. Color of bars illustrates percent incidence of major EAE subtype (classic: grey, or axial rotary (AR): orange). (**C**) Percent incidence of relapsing remitting (RR; green bars) and monophasic (light green bars) EAE in CC strains, with B6 mice shown as a reference control. (**D**) Comparison of EAE disease severity in CC strains, as calculated using cumulative disease score (see Materials and Methods), versus B6 reference controls. Significance of differences of each CC strain from B6 reference controls was determined via one-way ANOVA with Dunnett’s multiple comparisons test and indicated by asterisks where significant (* p<0.05, ** p<0.01, *** p<0.001, **** p< 0.0001), with corresponding bar colors indicating directionality as compared to B6, ie. blue = less severe and red = more severe. (**E-K**) Distribution of strain (**E**) cumulative disease score (CDS), (**F**) total EAE incidence, and incidence of (**G**) classic EAE, (**H**) AR-EAE, (**I**) chronic EAE, (**J**) RR-EAE, and (**K**) monophasic EAE, grouped by *H2^b^* and *H2^g7^* homozygous haplotypes. Each data point in panels (**E** - **K**) represents the average for a single strain. Significance of differences between haplotype groups was determined by unpaired T-test and indicated by asterisks where significant.

**Table 1.** Characteristics of CC strains used in EAE studies and their phenotypes

| CC Strain | Abbreviated Strain Name | H2 Haplotype | Founder Strain | n | Cohort (C#) | EAE Phenotype<sup>†</sup><br>Course (Subtype) | Sex Difference<sup>††</sup> |
| --- | --- | --- | --- | --- | --- | --- | --- |
| CC001/Unc | CC001 | b | B6 | 9 (4M, 5F) | C1 | Chronic (Classic) | None |
| CC002/Unc | CC002 | b | 129S1 | 11 (6M, 5F) | C1,C2, and C3 | Relapsing Remitting (Classic) | None |
| CC003/Unc | CC003 | b/g7 | 129S1/NOD | 10 (5M, 5F) | C1 and C2 | Monophasic (Classic) | None |
| CC004/TauUnc | CC004 | b | 129S1 | 14 (7M, 7F) | C1 and C2 | Severe Chronic (AR) | None |
| CC006/TauUnc | CC006 | b/g7 | B6/NOD | 10 (5M, 5F) | C1 and C2 | Mild Relapsing Remitting (Classic) | None |
| CC010/GeniUnc | CC010 | b | B6 | 10 (5M, 5F) | C1 and C2 | Chronic/Relapsing Remitting (Classic) | None |
| CC011/Unc | CC011 | g7 | NOD | 10 (5M, 5F) | C1 and C3 | Resistant (Classic) | None |
| CC018/Unc | CC019 | b | 129S1 | 9 (5M, 4F) | C1 and C3 | Chronic (Classic) | None |
| CC020/GeniUncJ | CC020 | b | B6 | 10 (5M, 5F) | C4 | Mild Monophasic (Classic) | None |
| CC023/GeniUnc | CC023 | b | 129S1 | 10 (5M, 5F) | C1 and C2 | Mild Chronic (Classic) | None |
| CC028/GeniUnc | CC028 | b | 129S1 | 10 (5M, 5F) | C1 and C3 | Severe Progressive (Classic) | M > F |
| CC030/GeniUnc | CC030 | g7/PWK | NOD/PWK | 10 (5M, 5F) | C1 and C2 | Chronic (Classic) | None |
| CC031/GeniUnc | CC031 | b | B6 | 9 (4M, 5F) | C3 | Mild Monophasic (Classic) | None |
| CC032/GeniUnc | CC032 | b | B6 | 10 (5M, 5F) | C1 and C2 | Mild Monophasic (Classic) | None |
| CC036/Unc | CC036 | g7 | NOD | 10 (4M,6F) | C1 and C2 | Mild Monophasic (Classic) | None |
| CC037/TauUnc | CC037 | b | B6 | 10 (5M, 5F) | C1 | Chronic/Relapsing Remitting (Classic) | None |
| CC038/GeniUnc | CC038 | b | 129S1 | 10 (5M, 5F) | C3 | Mild Chronic (AR) | M > F * |
| CC040/TauUnc | CC040 | b | B6 | 9 (4M, 5F) | C1 | Mild Chronic (Classic) | None |
| CC041/TauUnc | CC041 | b | 129S1 | 10 (5M, 5F) | C1 | Chronic (Classic) | None |
| CC042/GeniUnc | CC042 | b | 129S1 | 10 (5M, 5F) | C1 and C2 | Chronic (Classic) | M < F * |
| CC043/GeniUnc | CC043 | g7 | NOD | 10 (5M, 5F) | C1 and C2 | Secondary Progressive (Classic) | None |
| CC044/Unc | CC044 | b/g7 | B6/NOD | 10 (5M, 5F) | C1 and C2 | Mild Monophasic (Classic) | None |
| CC046/Unc | CC046 | g7 | NOD | 10 (5M, 5F) | C3 | Chronic (Classic) | M > F |
| CC051/TauUnc | CC051 | b | 129S1 | 10 (5M, 5F) | C1 | Relapsing Remitting (Classic) | None |
| CC059/TauUnc | CC059 | b/a | 129S1/A/J | 10 (5M, 5F) | C1 | Relapsing Remitting (Classic) | None |
| CC061/GeniUnc | CC061 | b | 129S1 | 11 (6M, 5F) | C1,C2, and C3 | Monophasic (AR & Classic) | None |
| CC068/GeniUnc | CC068 | b | 129S1 | 9 (4M, 5F) | C1 and C2 | Mild Monophasic (Classic) | None |
| CC072/GeniUnc | CC072 | b | B6 | 10 (5M, 5F) | C1,C2, and C3 | Mild Chronic (AR) | M < F * |
| CC074/Unc | CC074 | g7 | NOD | 10 (5M, 5F) | C1 and C2 | Mild Monophasic (Classic) | None |
| CC075/Unc | CC075 | b | 129S1 | 10 (5M, 5F) | C1 and C2 | Chronic (Classic) | None |
| CC083/Unc | CC083 | g7 | NOD | 10 (5M, 5F) | C1 | Chronic (AR & Classic) | None |
| CC084/TauUnc | CC084 | b/z | B6/NZO | 10 (5M, 5F) | C3 | Chronic (AR & Classic) | M < F * |

<sup>†</sup> EAE phenotype denotes severity (Mild, average- no title, and severe) as determined by cumulative disease score analysis compared to B6 reference controls (Figure 1A), disease course (chronic, relapsing remitting, or monophasic), as determined by greatest percent incidence except in the case of equal incidences which are shown with a /, and subtype (classic or AR), where AR EAE is classified by >25% AR-EAE incidence
<sup>††</sup>Sex difference reported as most significant P-value in any of the following: cumulative disease score, classic cumulative disease score, and axial rotary cumulative disease score
\*Denotes statistical significance (p<0.05) utilizing uncorrected Fisher's LSD

Male and female mice (n=5 of each sex), 9-14 weeks old, of each of the selected 32 CC strains, as well as B6 reference controls, were immunized subcutaneously with 200 µg MOG_35-55_ emulsified in CFA and received a single intraperitoneal injection of 200 ng pertussis toxin (PTX) as an ancillary adjuvant, both administered on day 0. Due to availability, CC mice were obtained and studied in 4 separate cohorts (**Table 1**), each of which included reference B6 controls. Mice were observed daily for a total of 50 days for the presence of clinical disease symptoms using the both the classical EAE^24^ and a modified axial rotary-EAE (AR-EAE)^43^ scoring scale as previously described (see Materials and Methods for a detailed description). Daily disease scores (**Fig. S2 – S7** and **Table S2 – S4**) were utilized to calculate EAE quantitative trait variables (QTVs) and derive EAE phenotypes (**Table 1** and **Table S5**) (see Materials and Methods).

In control B6 mice, this protocol resulted in a high penetrance (∼92%) of typical symptomatology manifesting as ascending paralysis (here termed “classic EAE”), with a moderately severe chronic disease phenotype (**Fig. 1B – D** and **Fig. 2A** and **B**). In contrast, the CC strains demonstrated a broad range of EAE phenotypes, with the overall incidence of EAE (of any type) ranging from 10-100% (**Fig. 1B**; **Table 1**, and **Table S5**). Analysis of disease course and EAE phenotypic traits (see Materials and Methods) classified CC strains into unique disease profiles/subtypes (**Table 1**). Besides classic EAE, a number of strains exhibited a high incidence of atypical AR-EAE, manifesting as severe ataxia and axial rotational body movements (**Fig. 1B**; **Table 1**, and **Table S5**). Additionally, a number of strains exhibited relapsing-remitting (RR)-EAE, or monophasic disease (**Fig. 1C**; **Table 1**, and **Table S5**). We used cumulative disease score (CDS; the total sum of daily scores accounting for both classic and AR-EAE subtypes) (see Materials and Methods) as a single quantitative variable capturing overall EAE severity/duration, which ranged greatly across the CC panel, with several strains demonstrating significantly lower CDS compared with B6, and two strains demonstrating significantly higher CDS (**Fig. 1D**; **Table 1**, and **Table S5**).

**Figure 2.**
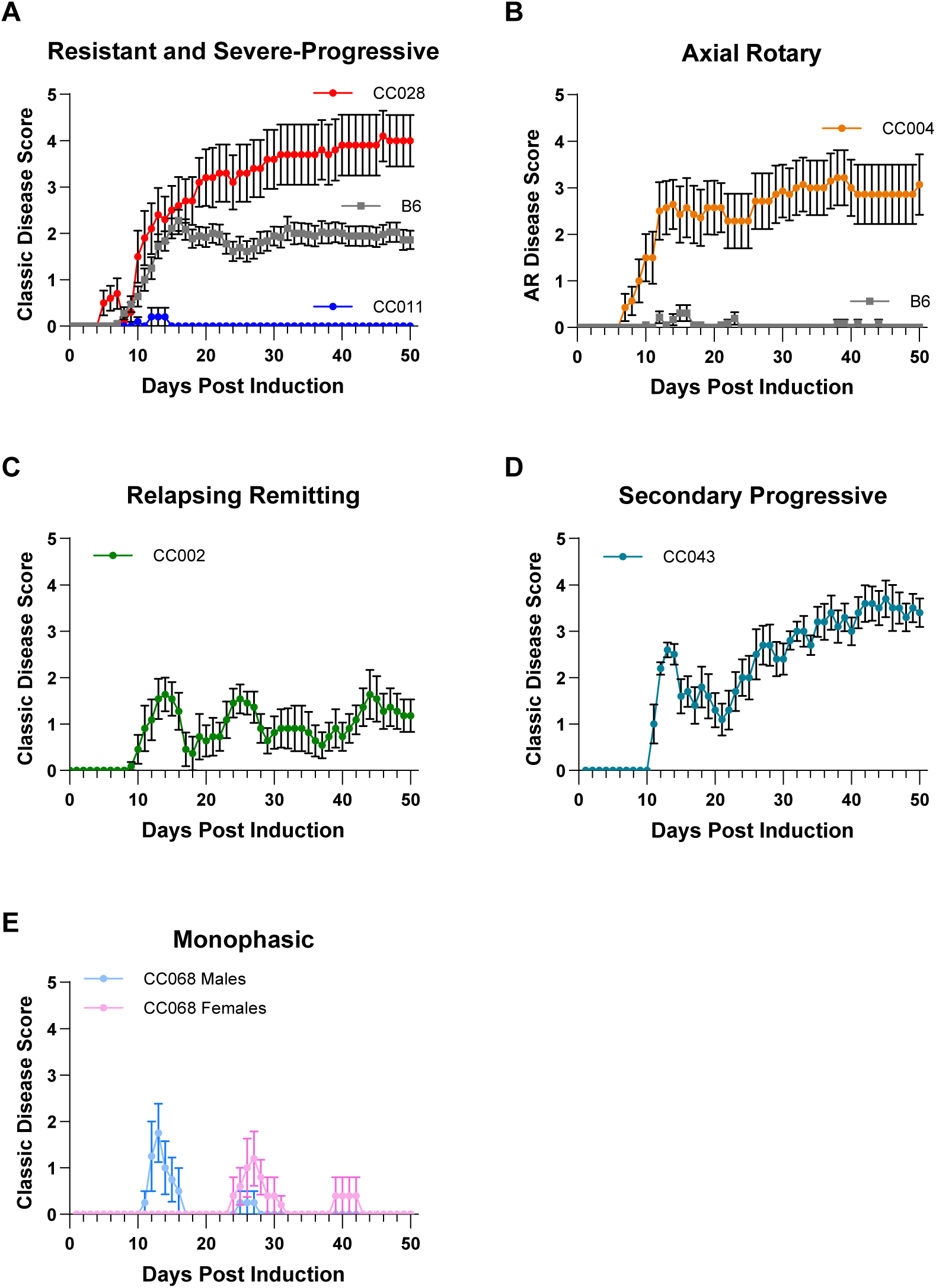
EAE in CC strains captures clinically relevant disease courses. EAE was induced and observed for 50 days in CC and B6 reference control mice as described in Figure 1. Post observation, disease course profiles for each strain were derived from daily disease scores (see Materials and Methods), revealing several strains that captured phenotypes of interest. These included: (**A**) severe progressive disease in CC028 (5M + 5F) mice (shown in red) and EAE resistance in CC011 (5M + 5F) mice (shown in blue), compared to B6 (18M + 18F) reference controls (shown in grey) (sexes pooled), (**B**) AR-EAE in CC004 (7M + 7F) mice (shown in orange) (sexes pooled), (**C**) relapsing remitting (RR)-EAE in CC002 (6M + 5F) mice (sexes pooled), (D) secondary progressive EAE in CC043 (5M + 5F) mice (sexes pooled), and (**E**) monophasic EAE in CC068 (4M + 5F) mice (sexes shown separately due to timing of disease onset). All panels show classic EAE scores, except (**B**), which shows AR-EAE scores, as indicated on the Y axis in each panel.

Because MHC class II alleles are the major genetic determinant of susceptibility to MS^10,11,47^, we next asked whether the limited *H2* haplotypes captured in our subset CC strains influenced EAE incidence and/or severity. Stratifying the CC strains by major *H2* haplotype (*H2^b^* or *H2^g7^*) demonstrated no significant difference in EAE cumulative disease score or incidence of any of the major EAE subtypes (**Fig. 1E – K**). Further subsetting the *H2* haplotype by founder strain of origin (B6, 129S1, or NOD) also did not reveal any significant differences (**Fig. S8**).

While many CC strains manifested clinical EAE presentation similar to B6, several strains captured extreme ends of the different phenotypes studied (**Fig. 2A-D**; **Table 1** and **Table S5**). These included diversity in both susceptibility and severity, from nearly completely resistant (CC011), to highly susceptible (CC028); with the latter presenting with rapidly progressing severe disease, with 70% (5/5 males and 2/5 females) reaching quadriplegia/humane endpoint by Day 39 (**Fig. 2A**). CC004 mice presented with the highest incidence (79%; **Fig. 1B**) of atypical AR-EAE with a severe chronic disease course (**Fig. 2B**), with a significantly higher overall CDS compared with B6 mice (**Fig. 1D**). Additionally, several strains exhibited diversity in disease course. Four strains exhibited a ≥ 50% incidence of RR-EAE, with CC002 being the most robust among them (**Fig. 1C** and **2C**). CC043 mice demonstrated a secondary progressive disease course (**Fig. 2D**). Several other strains, in particular CC068, presented with a monophasic disease course, which in the case of CC068 was staggered in day of onset by sex (**Fig. 1C** and **2E**). Taken together, these results demonstrate that the genetic diversity in CC mice captures a wide spectrum of clinically relevant EAE phenotypes, with several strains capturing extreme EAE phenotypes of interest for follow-up analysis.

MS exhibits well-documented sex differences in both disease incidence (higher in females) and disease progression (more severe in males)^6^. Sex differences in EAE have also been reported, predominantly in SJL/J mice and less so in B6 mice^48^. With our sample size (n=5 per sex per CC strain), while we were likely underpowered to detect subtle sex differences in individual strains, we could potentially capture larger effects if any were present. A two-way analysis of variance of the effect of strain and sex on CDS (analyzed separately for classic EAE, AR-EAE, or combined disease) revealed a highly significant effect of strain (as expected), no overall effect of sex, and a significant strain by sex interaction in the case of classic EAE CDS (**Fig. 3A-C**). A post-hoc analysis of the effect of sex on disease course within each CC strain identified significant and bi-directional effects of sex on EAE CDS in a total of 5 different strains across the different disease types (**Fig. 3A-C**, **Table 1**, and **Table S6**). Examination of disease course revealed distinct differences for classic EAE in CC046 (greater severity in males) (**Fig. 3D**) and CC042 (greater severity in females) (**Fig. 3E**), and for AR-EAE course in CC038 (greater severity in males) (**Fig. 3F**) and CC072 (greater severity in females) (**Fig. 3G**). Taken together, these results demonstrate that the effect of sex on EAE is highly genotype-dependent, indicating the presence of gene-by-sex interactions.

**Figure 3.**
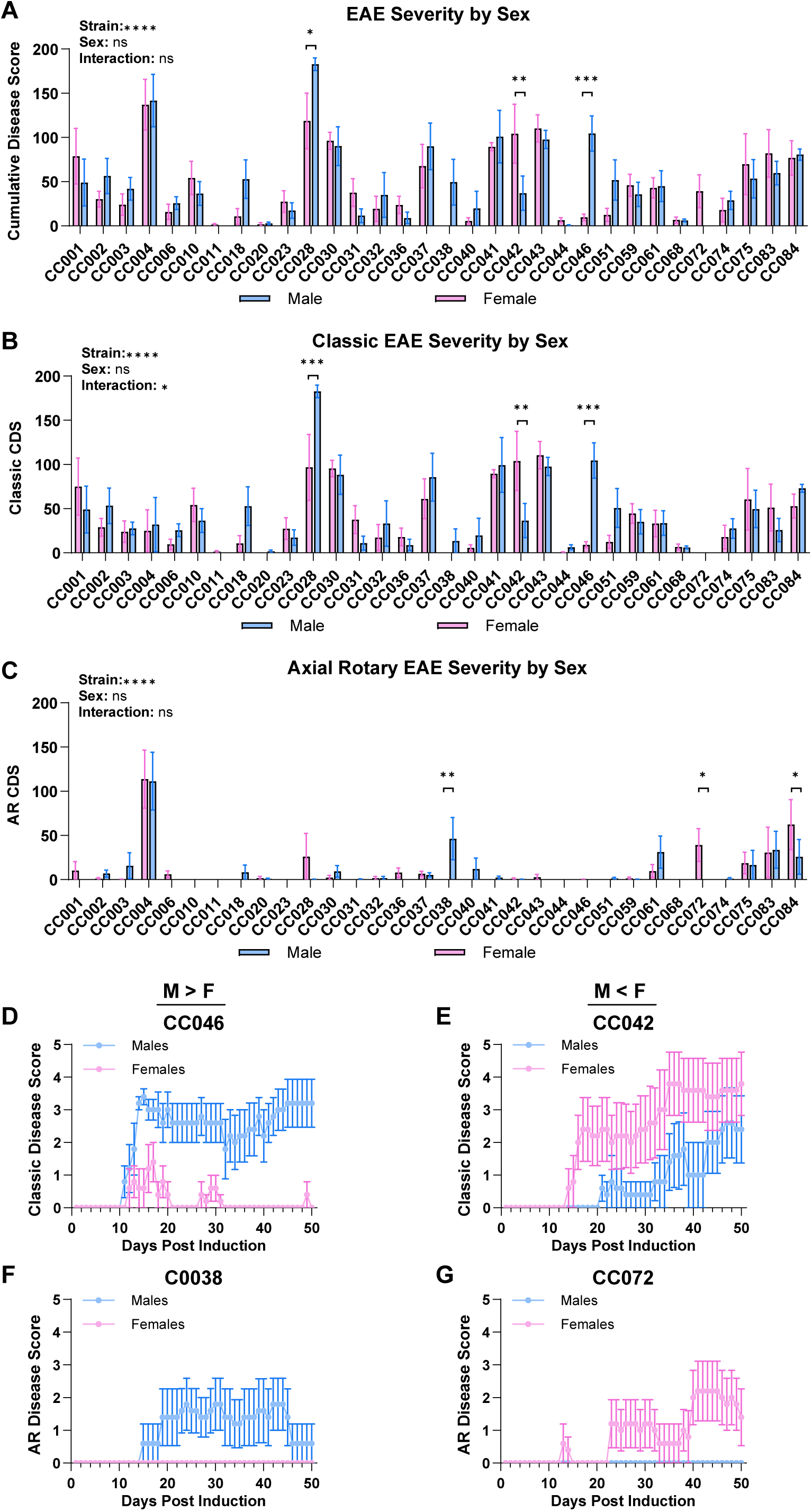
EAE in CC strains demonstrates bi-directional effects of sex on disease course. EAE was induced and observed for 50 days in CC mice as described in Figure 1. Disease severity was assessed for effects of sex within strain using (**A**) cumulative disease score (CDS; the total sum of disease scores), (**B**) classic cumulative disease score (classic-CDS; the total sum of classic disease scores), and (**C**) AR cumulative disease score (AR-CDS; the total sum of AR disease scores) (see Materials and Methods). Significance of differences was determined by two-way ANOVA and displayed with Fishers LSD multiple comparisons (see Table S6). Comparisons are indicated by asterisks where significant (* p<0.05, ** p<0.01, *** p<0.001, **** p< 0.0001). Disease course profiles of sex differences in classic EAE presentation in (**D**) CC046, and (**E**) CC042 mice, and AR-EAE presentation in (**F**) CC038, and (**G**) CC072 mice. Panels (**D**) and (**E**) display classic EAE scores while panels (**F**) and (**G**) display AR-EAE scores, as indicated on the Y axis in each panel.

### Distinct immunopathology in the spinal cord and brain is associated with classic and AR-EAE clinical phenotypes in CC028 and CC004 mice

To determine the immunopathological basis of the distinct EAE phenotypes identified in CC mice, we specifically focused on strains exhibiting RR-EAE (CC002), AR-EAE (CC004), and severe progressive EAE (CC028), together with reference B6 controls. In the experiments described above, brain and spinal cord were collected at D50 post EAE induction (or at time of humane endpoint euthanasia), fixed in formalin, and processed for sectioning and staining with hematoxylin and eosin (H&E) or Luxol fast blue (LFB) to assess immune cell infiltration and demyelination, respectively, using a semi-quantitative assessment by a blinded observer, similar to what we have previously published^49,50^ (see Materials and Methods). Analysis of spinal cord inflammation in B6 mice revealed expected focal inflammatory infiltrates, which were comparable in CC002 mice, and significantly reduced in CC004 mice (**Fig. 4A** and **B**). In contrast, spinal cords from CC028 mice demonstrated significantly greater levels of inflammation compared with B6 reference controls, characterized by extensive and dense infiltration of inflammatory cells (**Fig. 4A** and **B**). Surprisingly, the overall extent of demyelination in the spinal cord showed no significant difference in between the three CC strains and B6 reference control (**Fig. 4C** and **D**). Assessment of brain pathology revealed a CC004-specific increase in inflammation compared with B6 reference controls, characterized by prominent perivascular infiltrates and moderate to severe parenchymal infiltration in the cerebellum (**Fig. 4E** and **F**). Likewise, LFB-stained CC004 brain sections revealed a significant increase in the level of demyelination, defined by widespread white matter pallor affecting nearly all (>75%) of the sample (**Fig. 4G** and **H**). Taken together, these data demonstrate that AR-EAE clinical presentation in CC004 mice is associated with lesions in the cerebellum rather than the spinal cord, consistent with previous findings on AR-EAE^43,51^, while severe and rapidly progressing classical EAE in CC028 mice is associated with augmented spinal cord inflammation.

**Figure 4.**
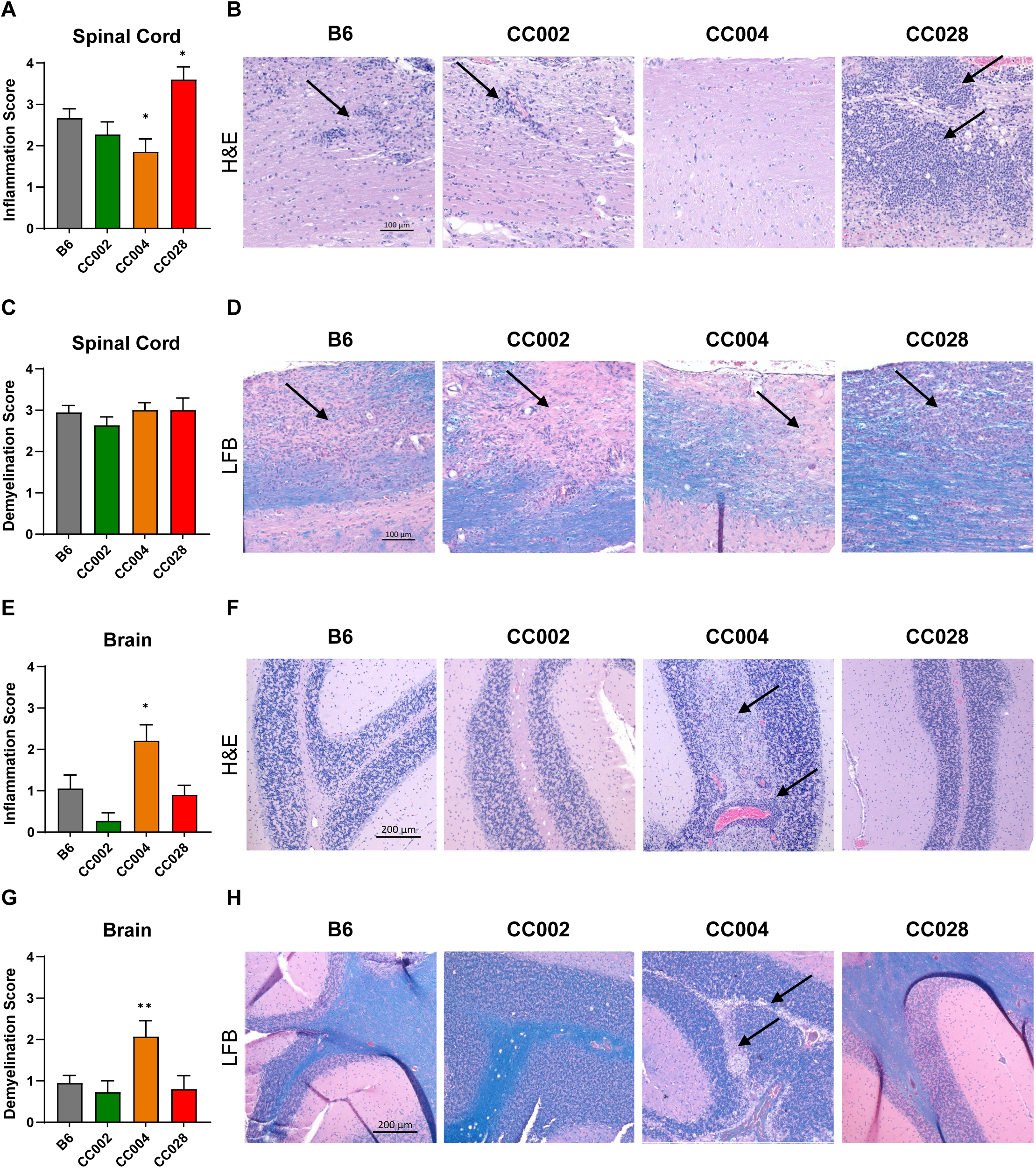
Severe progressive EAE in CC028 mice and AR-EAE in CC004 mice is associated with distinct pathology in the spinal cord and brain, respectively. EAE was induced and observed for a total of 50 days as described in Figure 1. At day 50, or at time of humane endpoint euthanasia, spinal cord and brains were collected and processed for sectioning and staining with H&E +/− LFB. Histopathologic evaluation of samples from B6 reference control (9M + 9F), CC002 (6M + 5M), CC004 (7M + 7F) and CC028 (5M + 5F) mice, sexes pooled, was performed as described in Materials and Methods. (**A**) Spinal cord inflammation scores, displayed as strain averages, and (**B**) corresponding representative spinal cord images for B6, CC002, CC004 and CC028 mice. Images were captured with a 10x objective, scale bar represents 100 μm, and arrows mark focal inflammatory infiltrates. (**C**) Spinal cord demyelination scores displayed as strain averages, and (**D**) corresponding representative spinal cord images for B6, CC002, CC004 and CC028 mice. Images captured at 10x objective, scale bar represents 100 μm, and arrows denote areas of demyelination. (**E**) Brain inflammation scores displayed as strain averages, and (**F**) corresponding representative brain images for B6, CC002, CC004 and CC028 mice. Images captured at 5x objective, scale bar represents 200 μm, and arrows mark regions of inflammatory infiltrates. (**G**) Brain demyelination scores displayed as strain averages, and (**H**) corresponding representative brain images for B6, CC002, CC004 and CC028 mice. Images captured at 5x objective, scale bar represents 200 μm, and arrows mark areas of demyelination. Significance of difference determined by ordinary one-way ANOVA, with Fishers LSD multiple comparisons to B6 reference controls, or by Brown-Forsythe and Welch ANOVA, with unpaired T test with Welch’s correction for multiple comparison testing when appropriate. Comparisons are indicated by asterisks where significant (* p<0.05, ** p<0.01, *** p<0.001, **** p< 0.0001).

### Severe EAE in CC028 and CC004 is associated with a greater abundance of myeloid rather than lymphoid cells in the CNS

To further characterize the extent of immune infiltration and immunopathology in severe-progressive and AR-EAE phenotypes in CC028 and CC004 mice, EAE was induced and followed for 14 days (D14) in order prevent CC004 and CC028 mice from succumbing to humane endpoints, while also capturing peak disease activity, with disease course recapitulating our previous results above (**Fig. 5A** and **B**). At D14 brain and spinal cord were collected and processed independently for leukocyte isolation, staining, and flow cytometric analysis (see Materials and Methods). Resulting cells were gated as shown in **Figure 5C** to assess key immune cell populations, including microglial cells (CD45^int^CD11b^+^CX3CR1^+^), myeloid cells (CD45^+^CD11b^+^Cx3CR1^low/-^), neutrophils (CD45^+^CD11b^+^CX3CR1^-^Ly6G^+^), B cells (CD45^+^CD11b^-^CD19^+^), and T cells (CD45^+^CD11b^-^CD19^-^ TCRβ^+^), as well CD4^+^ T cells producing the two signature cytokines, IFNγ and IL-17, for Th1 and Th17 cells, respectively. Assessment of these populations revealed an increase in total CD11b^+^ cell frequency in both the brain and spinal cord of CC004 and CC028 compared with B6 mice (**Fig. 5D** and **J**). Further examination of this population in the spinal cord revealed no significant differences in the population of microglia, myeloid cells, or neutrophils between the three strains/phenotypes (**Fig. 5E-G**). However, analysis of these same populations in the brain revealed a significantly greater population of microglia cells in CC028 mice compared with B6 (**Fig. 5K**). Additionally, we found a significant increase in myeloid cells in the brain of CC004 mice compared with B6 (**Fig. 5L**), which was mostly accounted for by an increase in neutrophils (**Fig. 5M**). The increased frequencies of myeloid cells in the brain and spinal cord of CC004 and CC028 mice compared with B6 mice were counterbalanced by a lower frequency of B and T cells, reported as frequency of CD45^+^ cells (**Fig. 5H-I, N-O**). A similar pattern was seen for IFNγ production by CD4 T cells, which was reduced in CC004 and CC028 compared with B6 mice, in both tissues (**Fig. 5P** and **R**). This decrease in Th1 cells was not accompanied by a reciprocal change in Th17 cells, as evidenced by lack of significant differences in IL-17 expression in CD4+ T cells (**Fig 5Q** and **S**). Taken together, these findings suggest that the severe classic EAE phenotype of CC028 mice is surprisingly associated with a decrease in lymphocyte abundance in the spinal cord, including greatly reduced Th1 cells. While a similar decrease in lymphocytes is seen in the brain and spinal cord of AR-EAE CC004 mice, there is an additional brain-specific increase in neutrophil infiltration, which is in alignment with the brain-specific increase in inflammation and demyelination observed during histopathological analysis (**Fig. 4**).

**Figure 5.**
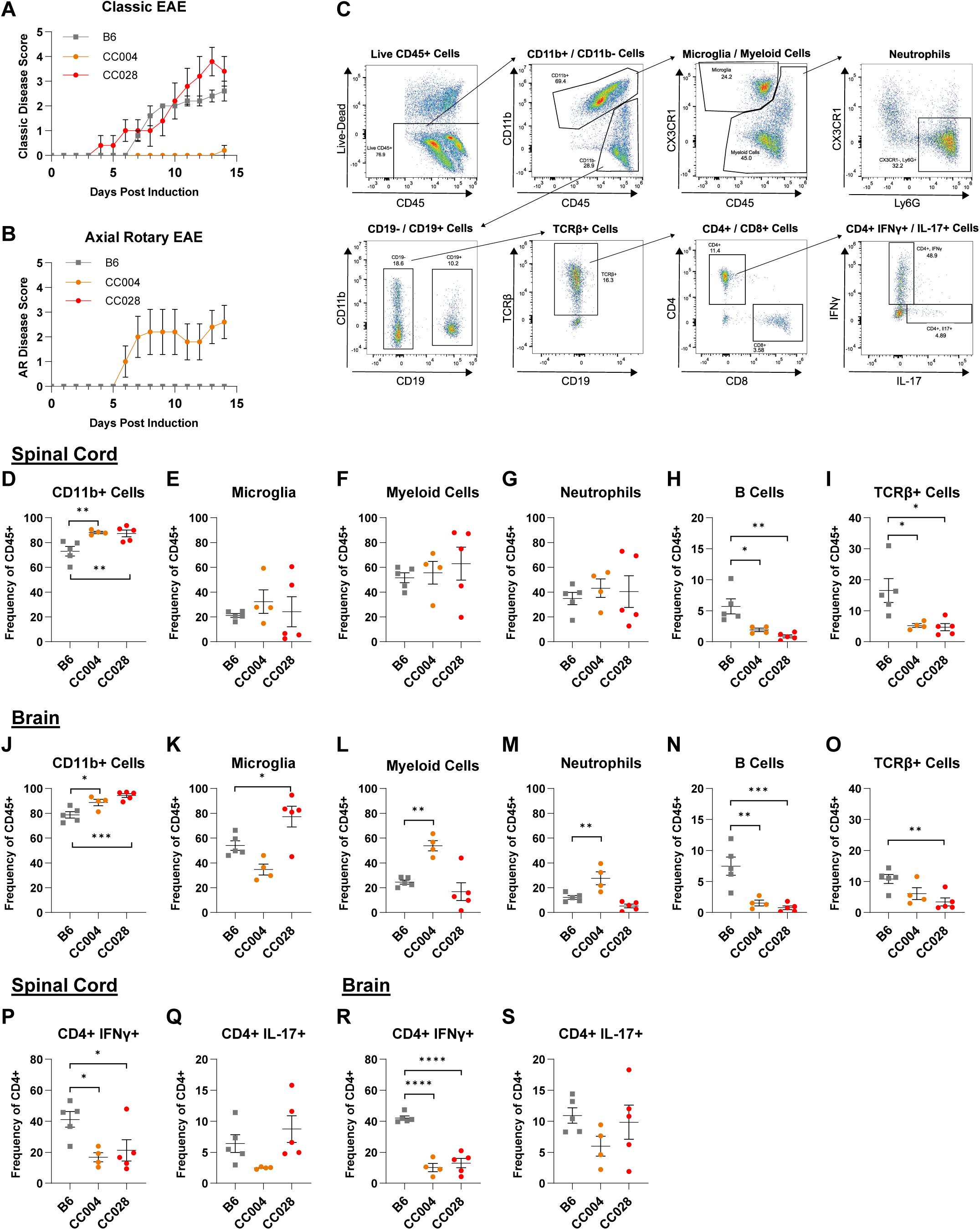
Severe EAE in CC028 and CC004 mice is associated with unique CNS immune profiles. EAE was induced in 8-13 week old male B6, CC004, and CC028 mice (5 per strain) by s.c. immunization with 200 µg of MOG_35-55_ emulsified in CFA, and a single i.p. injection of 200 ng of PTX on D0 (see Materials and Methods). Mice were observed daily for a total of 14 days to capture peak disease activity. On day 14 spinal cord and brains were collected and processed for flow cytometric staining (see Materials and Methods). Disease course profiles for B6, CC004, and CC028 mice displayed as (**A**) classic EAE or (**B**) AR-EAE. (**C**) Representative gating scheme for flow cytometric analysis. Scatterplots demonstrating key immune cell subsets in the spinal cord of B6, CC004 and CC028 mice, including (**D**) CD11b^+^ cells (CD45^+^CD11b^+^), (**E**) microglial cells (CD45^int^CD11b^+^CX3CR1^+^), (**F**) myeloid cells (CD45^+^CD11b^+^Cx3CR1^low/-^), (**G**) neutrophils (CD45^+^CD11b^+^CX3CR1^-^Ly6G^+^), (**H**) B cells (CD45^+^CD11b^-^CD19^+^), and (**I**) T cells (CD45^+^CD11b^-^CD19^-^TCRβ^+^). Scatterplots demonstrating key immune cell subsets in the brain of B6, CC004 and CC028 mice, including (**J**) CD11b^+^ cells, (**K**) microglial cells, (**L**) myeloid cells, (**M**) neutrophils, (**N**) B cells, and (**O**) T cells. Scatterplots demonstrating CD4^+^ T cells (CD45^+^CD11b^-^CD19^-^ TCRβ^+^CD4^+^) producing (**P**) IFNγ and (**Q**) IL-17 in the spinal cord of B6, CC004 and CC028 mice. Scatterplots demonstrating CD4^+^ T cells producing (**R**) IFNγ and (**S**) IL-17 in the brain of B6, CC004 and CC028 mice. Significance of differences of each CC strain from B6 reference controls was determined via one-way ANOVA with Dunnett’s multiple comparisons test and indicated by asterisks where significant (* p<0.05, ** p<0.01, *** p<0.001, **** p< 0.0001).

### Relapsing remitting EAE in CC002 mice is driven by both peripheral immune responses and non-hematopoietic derived factors

While peripheral immune cells initiate disease in EAE and MS, CNS-intrinsic factors play an important role in regulating disease progression. To determine which of these two distinct mechanisms serve as the basis for the genetically regulated relapsing remitting EAE phenotype in CC002 mice, reciprocal bone marrow chimera EAE experiments, between MHC-matched control B6 (*H2^b^*) and CC002 (*H2^b^*) mice were conducted (see Materials and Methods). In order to assess chimerism, we used congenic B6 mice carrying the CD45.1 allele (B6.SJL-Ptprca Pepcb/BoyJ), since we determined that CC002 mice carry a A/J-derived haplotype at the *Ptprc* locus, and therefore the CD45.2 allele. The donor/recipient combinations and respective CD45 alleles are detailed in **Figure 5A**, and we note that for the CC002→CC002 chimeras, congenic markers could not be used. At 8 weeks post bone marrow ablation and reconstitution, EAE was induced as above and observed for a total of 30 days, at which point spleen and CNS tissues were collected for immunophenotyping by flow cytometry (see Materials and Methods).

Flow cytometric analysis using CD45.2 and CD45.1 markers demonstrated successful bone marrow chimerism across all groups assessed, with some variation in chimerism across cell types as expected. An average of 95.4% (range 86.6 – 99.4%) of total splenic leukocytes were donor derived. For CD11b+ and CD19+ cells, the average chimerism was greater than 98% (**Fig. 6B** and **C**) cells. CD4^+^ and CD8^+^ T cells displayed an average of 81.2% and 78.2% chimerism, respectively, with increased to persistence of B6 host T cells in the CC002→B6 group (**Fig. 6D** and **E**). Subsequent assessment of donor cell frequency in the spinal cord demonstrated similar trends to those observed in the spleen, most notably the persistence of a significant fraction of host CD4^+^ and CD8^+^ T cells in the CC002→B6 group (**Fig. 6F** and **G**).

**Figure 6.**
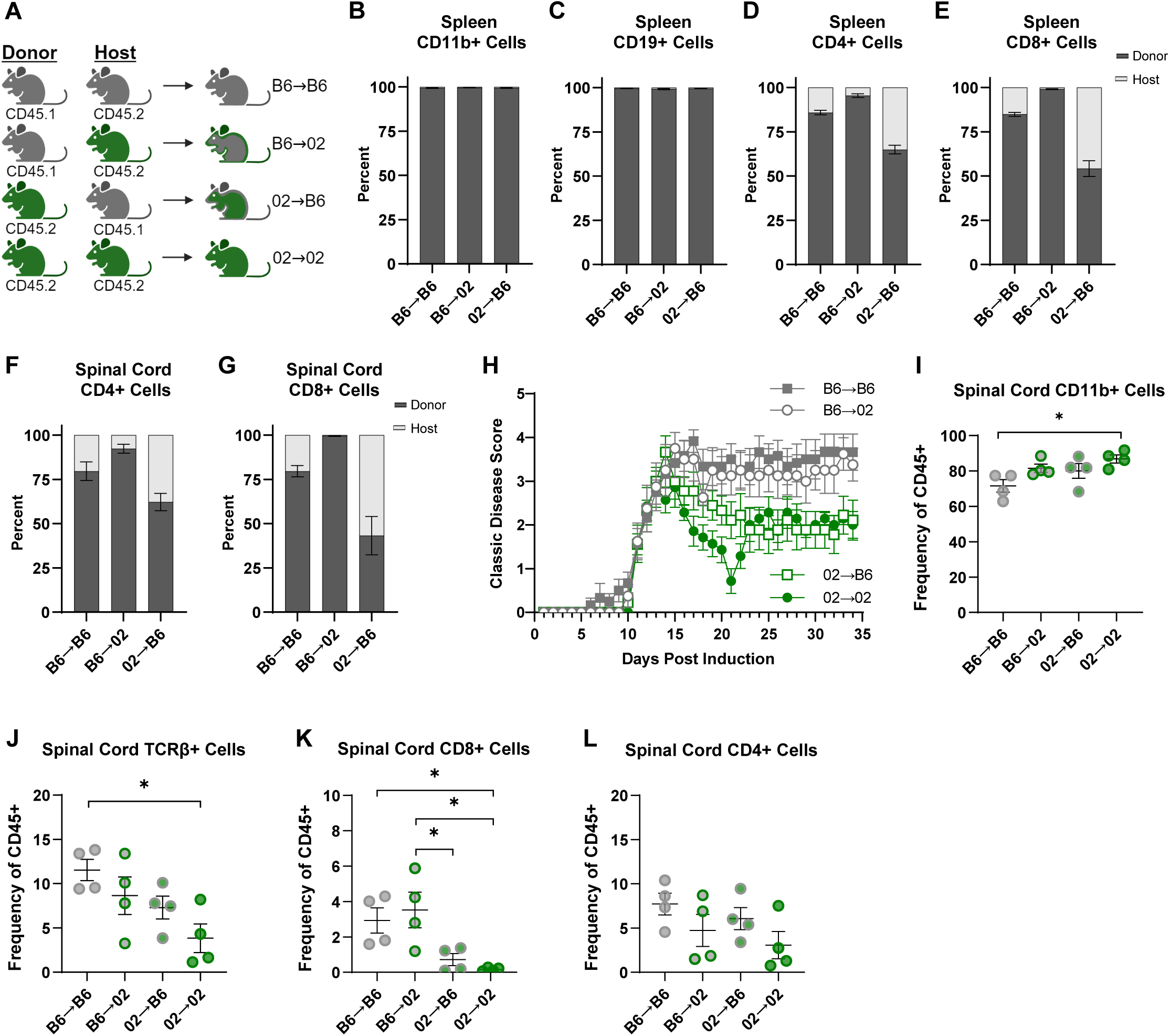
Peripheral immune and CNS intrinsic factors drive relapsing remitting EAE in CC002 mice. B6 and CC002 mice were subjected to bone marrow ablation and reconstitution to create reciprocal bone marrow chimeric mice (see Materials and Methods), designated as B6→B6 (7M + 4F), B6→02 (7M + 1F), 02→B6 (6M +4F), 02→02 (6M + 1F) and illustrated in the schematic in panel (**A**). Mice were rested for a total of 8 prior to EAE induction by s.c. immunization with 200 µg of MOG35-55 emulsified in CFA, and a single i.p. injection of 200 ng of PTX on D0 (see Materials and Methods). Mice were observed daily for a total of 34 days. On day 34 spleen, spinal cord and brain tissues were collected and processed for flow cytometric staining (see Materials and Methods). Percent chimerism was assessed in B6→B6, B6→02, and 02→B6 mice for splenic (**B**) CD11b^+^ cells (CD45^+^CD11b^+^CD19^-^), (**C**) CD19^+^ cells (CD45^+^CD11b^-^CD19^+^), (**D**) CD4^+^(CD45^+^CD11b^-^CD19^-^TCRβ^+^CD4^+^) and (**E**) CD8^+^ T cells (CD45^+^CD11b^-^CD19^-^TCRβ^+^CD8^+^), as well as infiltrating (**F**) CD4^+^ and (**G**) CD8^+^ T cells in the spinal cord. (**H**) Disease course profiles for B6→B6, B6→02, 02→B6, and 02→02 mice displayed as classic EAE. Comparison of spinal cord infiltrating immune cell populations in B6→B6, B6→02, 02→B6, and 02→02 mice for (**I**) CD11b^+^ cells, (**J**) CD19^+^ cells, (**K**) CD8^+^, and (**L**) CD4^+^ T cells. Significance of differences between groups was determined via one-way ANOVA with Tukey’s multiple comparisons test and indicated by brackets and asterisks where significant (* p<0.05, ** p<0.01, *** p<0.001, **** p< 0.0001).

Analysis of EAE course demonstrated that phenotypes for the control groups matched our original findings (**Fig. 2C**), with chronic EAE observed in B6→B6 chimeras and RR-EAE in CC002→CC002 chimeras (**Fig. 6H**), confirming that the bone marrow ablation and transplantation procedure did not alter these phenotypes. Analysis of the experimental groups demonstrated that bone marrow from B6 donors was sufficient to alter the disease course in CC002 hosts, resulting in a “B6-like” chronic EAE phenotype (**Fig. 6H**). Meanwhile, transfer of CC002 bone marrow into B6 hosts resulted in remitting EAE, although without relapse (**Fig. 6H**). Assessment of CNS infiltrating cells by flow cytometry revealed a significant difference between CC002 and B6 mice with a greater frequency of CD11b^+^ cells (**Fig. 6I**) and a reduced population of TCRβ^+^ cells (**Fig. 6J**) in CC002 mice compared with B6. This reduction in TCRβ^+^ cells was driven primarily by a significant reduction in frequency of CD8^+^ cells in the strains with remitting EAE phenotypes (CC002→B6 and CC002→CC002) compared to chronic EAE phenotypes (B6→B6 and B6→CC002) (**Fig. 6K**), an effect that was not observed in the CD4^+^ cell population (**Fig. 6L**). Taken together, these results suggest that while remission of EAE in CC002 mice is intrinsic to peripheral immune system, CNS-intrinsic genetic factors may drive relapse. Alternatively, the lack of relapse in CC002→B6 chimeras could be driven by the significant percentage of remaining host (B6) T cells.

### QTL mapping reveals distinct loci associated with EAE subtype, course, and severity

Besides the identification of novel phenotypes, an advantage of the CC model is the ability to genetically map loci controlling specific phenotypic traits of interest, with mapping power and resolution increasing with the number of unique strains studied. We performed genome-wide association mapping for two major EAE quantitative traits: 1) EAE incidence (**Fig. 1B**), as a measure of disease susceptibility, and 2) cumulative disease score (CDS; **Fig. 1D**), as a cumulative measure of disease severity, duration, and incidence. For incidence, we either measured total EAE incidence, or subsetted incidence by: 1) disease subtype (classic EAE and AR-EAE), and 2) disease course (chronic, RR-EAE, and monophasic EAE). Association mapping was performed using R/qtl2 software^52^, with genome-wide significance logarithm of odds (LOD) thresholds determined by permutation analysis, using a relaxed threshold (given the limited number of strains studied and the complexity of the traits) of 20% to identify top QTL (see Materials and Methods). QTL were designated *Eaecc* (*EAE* QTL identified in *CC* mice) and numbered in the order of their description in the manuscript (*Eaecc1-6*).

The first trait mapped was total EAE incidence (any disease subtype). While no single QTL passed the significance threshold, the lead QTL on proximal Chr4 (*Eaecc1*; LOD score = 6.79) fell just short of significance (**Fig. S9A** and **B; Table 2**). Subsetting EAE incidence by disease subtype and disease course revealed distinct linkage patterns for each classification and identified several QTL, some of which passed the significance threshold. While analysis of classic EAE (i.e. non-AR) incidence yielded a top QTL on Chr5 that fell short of significance (*Eaecc2;* LOD score = 6.64; **Fig. S9C** and **D; Table 2**), AR-EAE incidence mapping revealed a lead QTL on Chr9 (LOD score = 7.96) that passed 20% genome-wide significance (*Eaecc3;* **Fig. 7A**; **Table 2**). Closer examination of the founder effects at this QTL revealed that WSB, and to a lesser extent NZO, alleles were associated with higher incidence of AR-EAE, while NOD alleles were associated with lower incidence (**Fig. 7B**). Consistent with this, analysis of genotype by phenotype distribution revealed that of the 32 studied CC strains, the top five CC strains with the highest AR-EAE incidence (CC004, CC083, CC084, CC072, and CC038) carried either WSB or NZO alleles at the QTL region, aligning with the observed QTL founder effects (**Fig. 7C**). Subsetting EAE incidence by disease course revealed no major associations for chronic EAE (**Fig. S9E**), but RR-EAE and monophasic EAE revealed one QTL each on Chr18 (*Eaecc4;* LOD score = 6.40) and Chr6 (*Eaecc5;* LOD score = 7.31), respectively, that fell just short of the significance threshold (**Fig. S9F - I; Table 2**). Taken together, these results reveal distinct linkage patterns for the incidence of different EAE subtypes or disease courses, suggesting that these phenotypes are controlled by several distinct major loci, further supporting the idea that disease course in MS is genetically controlled.

**Figure 7.**
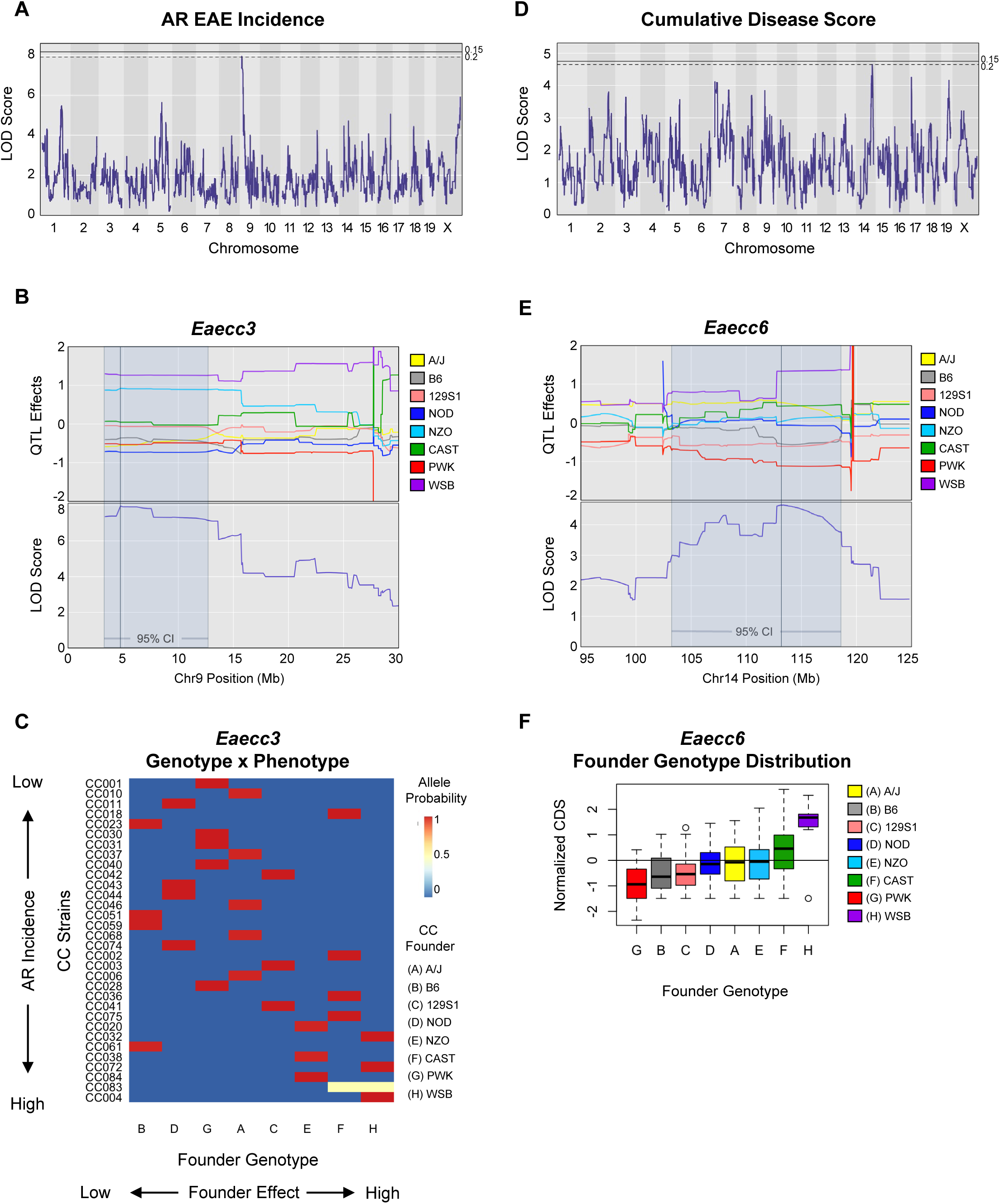
QTL analysis reveals distinct genetic linkage patterns for AR-EAE incidence and EAE severity. EAE was induced and evaluated in 32 CC strains, as described in Figure 1. EAE disease phenotypes and quantitative trait variables were calculated, and quantitative trait loci (QTL) mapping was performed (see Materials and Methods). (**A**) Manhattan plot demonstrating logarithm of odds (LOD) traces for QTL mapping assessing genome association with AR-EAE incidence. Genome wide significance thresholds of 15% (solid line) and 20% (dashed line) were determined by permutations (n=1000). (**B**) Corresponding CC founder allele effects plot for lead QTL identified on chromosome 9 - *Eaecc3*. (**C**) Heatmap demonstrating CC strain distribution based on genotype by phenotype analysis for the chromosome 9 AR-EAE incidence QTL – *Eaecc3*. (**D**) Manhattan plot demonstrating LOD traces for QTL mapping assessing genome association with EAE severity as determined by cumulative disease score. Permutation was used to determine Genome wide significance thresholds of 15% (solid line) and 20% (dashed line) were determined by permutations (n=1000). (**E**) Corresponding CC founder allele effects plot for lead QTL identified on chromosome 14 – *Eaecc6*. (**F**) Box and whisker plot demonstrating distribution of CC founder alleles within studied CC strains. Panels (**B**), (**C**), (**E**), and (**F**) use the conventional letter and color designations for CC founder strains as follows: A/yellow: A/J, B/grey: C57BL/6J, C/pink: 129S1/SvlmJ, D/dark blue: NOD/ShiLtJ, E/light blue: NZO/HILtJ, F/green: CAST/EiJ, G/red: PWK/PhJ, and H/purple: WSB/EiJ.

**Table 2.** Major EAE phenotype QTL detected in the CC

| QTV | QTL Symbol | Chr: Position (95% Confidence Interval) (Mb) | 20% Genome Wide Significance LOD Score | LOD Score | Top 5 Prioritized Genes FPR≥0.05 |  |
| --- | --- | --- | --- | --- | --- | --- |
|  |  |  |  |  | Lead Candidate Genes (Immune) | Lead Candidate Genes (CNS) |
| Total EAE Incidence | *Eaecc1* | 4: 4.33 (4.04-11.82) | 7.31 | 6.79 | *Trp53inp1, Plekhf2, Tox, Gem, Chd7* | *Trp53inp1, Tox, Sdcbp, Plekhf2* |
| Classic EAE Incidence | *Eaecc2* | 5: 74.06 (51.50-75.92) | 7.50 | 6.64 | *Rbpj, Klf3, Stim2, Fam114a1, Pcdh7* | *Klf3, Rasl11b, Apbb2, Atp8a1, Stim2* |
| AR-EAE Incidence | *Eaecc3* | 9: 4.75 (3.37-12.50) | 7.91 | 7.96 | *Yap1, Kbtbd3* | *Dync2h1, Birc3* |
| RR-EAE Incidence | *Eaecc4* | 18: 57.43 (56.12-60.12) | 7.20 | 6.40 | - | *Fbn2, Isoc1* |
| Monophasic EAE Incidence | *Eaecc5* | 6: 120.17 (118.97-121.49) | 7.65 | 7.31 | *Il17ra, Wnk1* | *Wnk1, Il17ra, Cecr2* |
| Cumulative Disease Score | *Eaecc6* | 14: 113.22 (103.97-118.63) | 4.65 | 4.65 | *Abbc4* | *Abbc4, Gpc6* |

Mapping of total EAE severity (encompassing both classic and AR-EAE presentation), using CDS as the quantitative trait, revealed a distinct linkage pattern, with the top associated QTL on distal Chr14 passing 20% genome-wide significance (*Eaecc6*; LOD score = 4.65; 20% threshold LOD score = 4.65) (**Fig. 7D**; **Table 2**). Closer examination of the founder effects at this QTL revealed that WSB alleles were associated with more severe EAE, while PWK (and to a lesser extent 129S1 and B6) alleles were protective (**Fig. 7E** and **F**). Taken together, these results reveal a novel QTL controlling EAE severity in CC mice, supporting the notion that disease severity in MS is a genetically regulated trait.

### Candidate gene prioritization nominates candidate genes for QTL controlling EAE traits

The ultimate goal of genetic mapping is to identify the causative genes underlying the associated phenotypes of interest, which, in recombinant inbred strain populations like the CC is often impeded by statistical resolution due to large haplotype blocks. This limitation can be overcome by gene prioritization approaches^53–57^. Our prioritization analysis focused on the two QTL reaching 20% genome wide significance: AR-EAE incidence (Chr9; 95% confidence interval ∼9.1Mb) and EAE severity (Chr14; 95% confidence interval ∼14.7Mb), as well as the suggestive narrow QTL associated with incidence of monophasic EAE (Chr6; 95% confidence interval ∼2.5Mb) (**Table 2**). While these QTL are moderately high resolution, these loci still contain numerous genes: Chr9 (81 genes, 36 protein-coding), Chr14 (130 genes, 19 protein-coding), and Chr6 (68 genes, 32 protein-coding). To prioritize lead candidate genes in an unbiased manner, we utilized a machine learning-based approach that we developed and described previously^55–57^, depicted schematically in **Figure 8A**. Briefly, this approach trains support vector machine (SVM) classifiers to distinguish trait-associated genes from randomly drawn genes using feature vectors derived from tissue-specific connectivity networks (functional networks of tissues in mouse (FNTM) database)^58^. The trained models are then asked to classify each positional candidate gene as trait-related or not trait-related. Here, we used the top 500 genes (as determined by -log10(p value)) associated with MS from the National Human Genome Research Institute GWAS catalog^59^ as the training set. We derived feature vectors for training from two tissue-specific mouse gene interaction networks from FNTM: 1) the immune system, the initiator and driver of pathology in EAE and MS) and 2) the CNS, as the target organ in EAE and MS (see Materials and Methods; **Table S7**).

**Figure 8.**
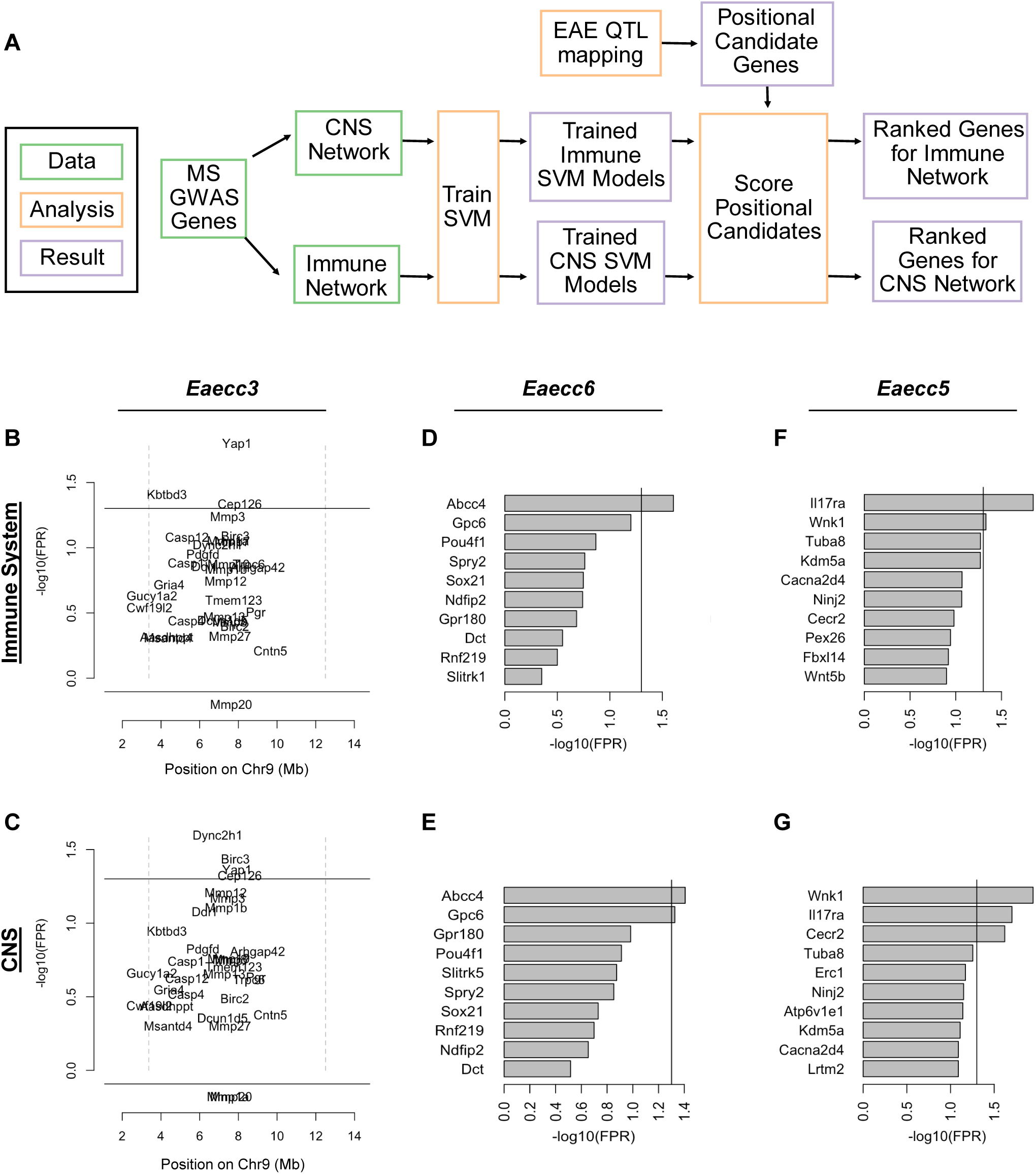
Machine learning-based functional candidate gene prioritization nominates distinct genes associated with QTL for AR-EAE incidence, EAE severity, and monophasic EAE incidence. Support vector machine (SVM) classifiers were trained using MS GWAS genes and integrated with extensive tissue specific connectivity networks for the CNS and immune system to rank gene candidates associated EAE QTL in the context of either the CNS or immune system (see Materials and Methods), as illustrated by the schematic in panel (**A**). Ranked candidate genes for the AR-EAE incidence QTL on chromosome 9 – *Eaecc3* for the (**B**) immune system network and (**C**) CNS network, genes are plotted by genomic position on the x axis and - log(FPR) on the y axis with the dotted lines demonstrating QTL boundaries. Ranked candidate genes graphed independent of genomic position for the cumulative disease score QTL on chromosome 14 – *Eaecc6* for the (**D**) immune system network and (**E**) CNS network, and the monophasic EAE incidence QTL on chromosome 6 – *Eaecc5* for the (**F**) immune system network and (**G**) CNS network. The solid line in panels **B** – **G** corresponds to the FPR threshold of 0.05.

For the QTL associated with AR-EAE incidence on Chr9, this approach prioritized a total of four genes (two immune system-specific and two CNS-specific) passing the false positive rate (FPR) cutoff at 0.05, with the two top prioritized genes being: *Yap1* (immune specific) and *Dync2h1* (CNS-specific) (**Fig. 8B** and **C**; **Table 2**). For the EAE severity QTL on Chr14, this approach prioritized two genes passing the FPR cutoff of 0.05: *Gpc6* (CNS-specific) and *Abcc4* (both tissues) (**Fig. 8D** and **E**; **Table 2**). Additionally, *Gpc5*, which is located adjacent to *Gpc6* within the Crh14 QTL, is itself an MS-GWAS candidate, although it was not ranked utilizing the above-described approach, due to insufficient connectivity in the networks. Assessment of the narrow QTL on Chr6 associated with monophasic EAE incidence prioritized a total of three genes (one CNS-specific and two in both tissues) passing the threshold of 0.05 FPR, with the top prioritized genes in the immune system and CNS identified as *Il17ra* and *Wnk1*, respectively, and both genes passing the FPR cutoff of 0.05 in both tissues (**Fig. 8F** and **G**; **Table 2**). Gene prioritization for the remaining suggestive QTL, including Chr4 total EAE incidence, Chr5 classic EAE incidence, and Chr18 RR-EAE incidence, identified additional lead candidate genes, including *Tox*, *Klf3*, and *Fbn2*, respectively (**Fig. S10** and **Table 2**). Taken together, this analysis prioritizes several genes as plausible candidates driving the EAE phenotypes of interest via effects on the immune system or the CNS.

Because missense variants have a high potential impact on gene function, we identified nonsynonymous single nucleotide variants (nsSNPs) distinguishing the founder alleles exerting the strongest opposing effects at each of the three lead QTL (**Fig. 7B** and **D; Fig. S9G**), focusing only on the top prioritized candidate genes (**Fig. 7**). Assessment of nsSNPs segregating between WSB (high AR-EAE incidence) and NOD (low AR-EAE incidence) at the top two prioritized genes (*Yap1* and *Dync2h1*) found no divergent nsSNPs. However, analysis of the remaining genes passing the 0.05 FPR cutoff revealed one nsSNP matching the strain distribution in *Birc3* (**Table 3**). For the Chr14 QTL controlling EAE severity, comparison of nsSNPs differentiating WSB (high EAE CDS) from PWK (low EAE CDS) at the top two prioritized genes revealed one nsSNP in *Gpc6*, and zero in *Abcc4* (**Table 3**). For the Chr6 QTL controlling monophasic EAE incidence, comparison of PWK (high monophasic EAE incidence) vs. 129S1 (low monophasic EAE incidence) alleles in *Il17ra* revealed no segregating nsSNPs, while two nsSNPs were identified in *Wnk1* (**Table 3**). Taken together, these results, combined with our prioritization analysis above, highlight potential coding variants driving the EAE phenotypes of interest, to be functionally validated in future studies, although we also do not discount the importance of non-coding regulatory variants.

**Table 3.**
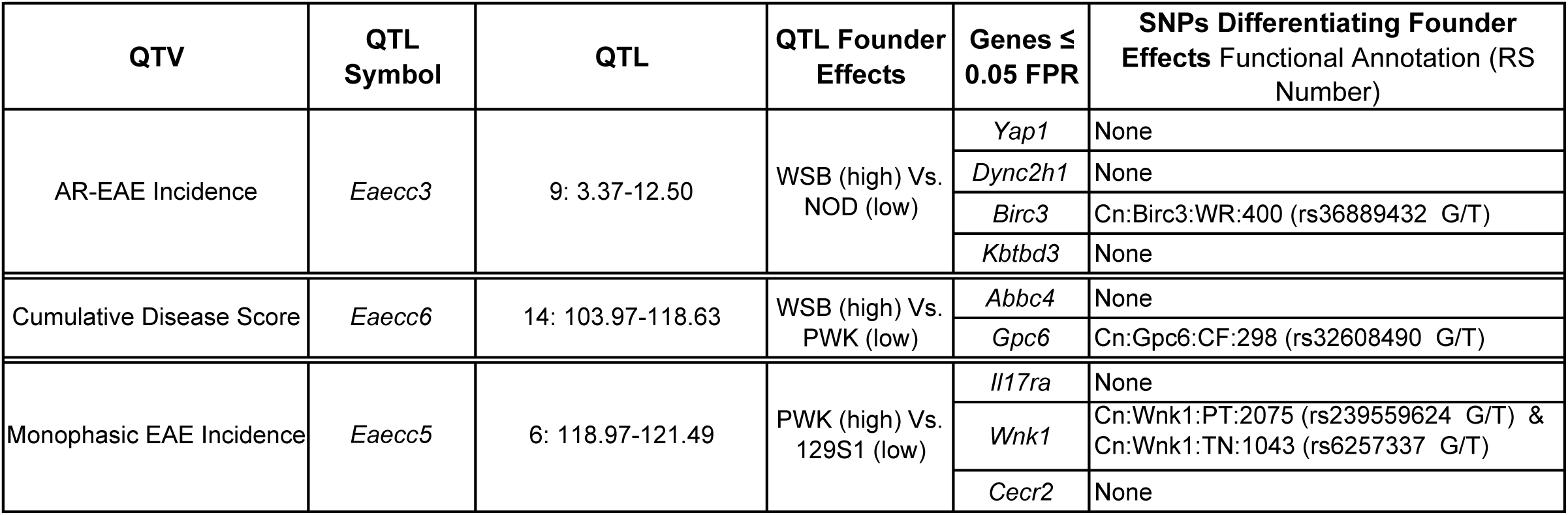
Coding non-synonymous variants in top prioritized QTL candidate genes

| QTV | QTL Symbol | QTL | QTL Founder Effects | Genes ≤ 0.05 FPR | SNPs Differentiating Founder Effects Functional Annotation (RS Number) |
| --- | --- | --- | --- | --- | --- |
| AR-EAE Incidence | Eaecc3 | 9: 3.37-12.50 | WSB (high) Vs. NOD (low) | Yap1 | None |
|  |  |  |  | Dync2h1 | None |
|  |  |  |  | Birc3 | Cn:Birc3:WR:400 (rs36889432 G/T) |
|  |  |  |  | Kbtbd3 | None |
| Cumulative Disease Score | Eaecc6 | 14: 103.97-118.63 | WSB (high) Vs. PWK (low) | Abbc4 | None |
|  |  |  |  | Gpc6 | Cn:Gpc6:CF:298 (rs32608490 G/T) |
| Monophasic EAE Incidence | Eaecc5 | 6: 118.97-121.49 | PWK (high) Vs. 129S1 (low) | Il17ra | None |
|  |  |  |  | Wnk1 | Cn:Wnk1:PT:2075 (rs239559624 G/T) & Cn:Wnk1:TN:1043 (rs6257337 G/T) |
|  |  |  |  | Cecr2 | None |

## Discussion

The genetic basis of disease course and/or pathology in MS remains obscure. To bridge the gap between genetic association studies and the cellular underpinnings of MS, animal models are needed, but most current mouse models fail to address the importance of natural genetic heterogeneity that is so pronounced in human populations. To expand the genetic diversity of the standard EAE model, we leveraged the CC - a highly genetically diverse mouse strain panel. This approach revealed a wide spectrum of distinct EAE phenotypes in CC mice. Immunological profiling of CC strains with clinically relevant phenotypes revealed distinct pathology associated with different EAE profiles. Additionally, QTL mapping and machine learning-based functional candidate gene prioritization revealed distinct genetic linkage patterns and identified several top candidate genes associated with the different EAE phenotypes captured by the CC mice. Taken together, these results demonstrate that introducing natural genetic diversity into the standard EAE model captures a wide spectrum of clinically relevant EAE phenotypes, providing a novel resource for the research community. Importantly, our results also provide support for the hypothesis that heterogeneity in disease course in MS has a strong genetic component.

The major risk genes for MS incidence reside in the MHC locus^9,11^. While we originally intended to use the full set of six MHC haplotypes represented across the CC population, this was precluded by the low efficacy of the “MHC-agnostic” EAE induction using mSCH (**Fig. S1**). Our MOG peptide-based EAE induction approach, while more efficient, cost-effective, and reproducible, allowed the inclusion of only two different compatible MHC haplotypes, *H2^b^* from 129S1 and B6 founder strains, and *H2^g7^* from NOD. While we found no significant effect of these two specific haplotypes on EAE phenotypes of interest (**Fig. 1E-K** and **Fig. S8**), this does not rule out a major role of additional MHC alleles in disease progression, which has been suggested in a several small association studies in humans^47,60^. This can be addressed in future studies using immunization with recombinant encephalitogenic proteins, such as MOG. Another approach could be to generate intercrosses between the *H2*-compatible CC strains in this study and additional CC strains carrying other *H2* haplotypes, to create heterozygosity at the MHC locus, a situation more representative of human genetics. Nonetheless, our approach, by limiting the MHC diversity, may have in fact increased our power to detect non-MHC genetic effects across the rest of the genome.

Relevant to the expansion of the EAE repertoire, this study represents a novel application of the widely used MOG_35-55_ EAE induction protocol to achieve a variety of disease course profiles across a panel of genetically distinct mice. Historically, the most common approach to reproducibly modeling a variety of distinct EAE disease profiles has been to utilize varying myelin derived peptides for EAE induction in corresponding susceptible strains of mice^61–63^. For example, chronic EAE disease profiles have been most commonly modeled using MOG_35-55_ induced EAE in B6 mice, while proteolipid protein (PLP) peptide 135-151-induced EAE in SJL/J (*H2^s^*) mice has been commonly used to model RR-EAE^61^. However, with the methods presented here, we were able to model a variety of distinct EAE disease profiles, including both chronic disease as well as RR disease, such as seen in CC002, using a single peptide induction approach, and in many cases on the same *H2* background. As such, the CC mice studied represent novel models for various MS disease courses for which adequate model representation has been somewhat limited. These include but are not limited to: CC068, with monophasic EAE as a potential model for clinically isolated syndrome; CC043, as a model for secondary progressive (SP)-MS, given the presence of a relapsing remitting phase that transitions into a progressive increase in clinical severity; CC028, as a potential model for primary progressive (PP)-MS, given the rapid onset and progressive increase in severity of disease; and CC004 with AR-EAE, as a model capturing ataxic symptomology of MS and brain lesions.

The occurrence of AR-EAE has been previously documented in several models, although the incidence varies and depends on the induction method. Greer and colleagues found a high penetrance of AR-EAE in PLP_109-209_ immunized C3H/HeJ mice (*H2^k^*), where it is was accompanied by distinct neuropathology, involving predominantly the cerebellum or brain stem^51^, consistent with our own findings. Follow-up studies from the Goverman group demonstrated that recombinant MOG-induced EAE in the closely related C3H/HeB (*H2^k^*) strain results predominantly in AR disease, whereas C3H.SW MHC congenic mice (carrying *H2^b^*) lose this phenotype and develop classic EAE, suggesting that AR-EAE incidence in this model is MHC-dependent^64^. Interestingly, they also found that AR-EAE was associated with higher Th17:Th1 ratios and could be induced more readily by adoptive transfer of Th17 cells. Consistent with this, several knockout strains on the B6 background, such as IFN-γ or *Socs3*-deficient mice, also show increased incidence of AR-EAE and brain pathology, which is often associated with increased neutrophil infiltration^43,65–68^. In our own studies, we found a similar increase in brain pathology in CC004 mice, which exhibited predominantly AR-EAE (**Fig. 3**). However, while we also found a brain-specific increase in neutrophil infiltration and a pronounced reduction in T helper IFNγ production in CC004 mice compared with B6, we did not find a corresponding increase in IL-17 production, which was in fact reduced by about two fold, albeit non-significantly (**Fig. 4**), suggesting that an increased Th17:Th1 ratio is not likely to be a major driver of AR-EAE in CC004 mice. Importantly, our QTL mapping results identified a non-MHC linked locus driving AR-EAE (*Eaacc3*) on Chr9, implicating novel genes in this phenotype (discussed below). Future mechanistic studies can further dissect whether the AR-EAE phenotype in CC004 mice is immunologically driven, and if so, whether neutrophils and/or Th17 cells play a major role. Importantly, cerebellar pathology and ataxia are a common feature of MS, whereas longitudinally extensive spinal cord pathology, while a major feature of “classic” EAE models, is less common in MS^69,70^.

Other distinct models of EAE disease progression in laboratory inbred strains of mice have been previously described. In particular, secondary progressive disease has been described in MOG_35-55_ induced NOD mice^71^ and mSCH-induced EAE in Biozzi ABH mice^72^. Much debate continues as to whether these models truly represent progressive disease representative of MS^44^, which has more neurodegenerative and less inflammatory characteristics^5^. While we identified two CC strains with either rapidly progressive severe disease (CC028) or secondary progression following a remission (CC043), we acknowledge that it is as yet unclear whether these represent valid models of progressive MS. In fact, compared with B6 mice, we observed greater histologic evidence of spinal cord inflammation in CC028 at early time points in disease (**Fig. 4**), suggesting a more severe inflammatory disease, although flow cytometric analysis also revealed reduced lymphocyte frequencies and increased frequencies of CD11b+ cells. Future studies focused on later time points can help to clarify the immunopathologic basis of progressive EAE in CC028 and CC043 mice.

In MS, sex differences in incidence (greater in females) and severity (greater in males) have been well documented^6,7^, but there is limited evidence for gene-by-sex interactions^73^, and GWAS studies typically use pooled male and female data to maximize power^9^. In contrast, the EAE model allows one to directly assess the effect of sex on the same fixed genotype, i.e. strain. While no uniform effect of sex was seen across all CC strains when combined, bi-directional sex differences were noted in several individual CC strains. Sex differences in EAE incidence were noted in CC038 (60% incidence in males and 0% incidence in females) and CC072 (0% incidence in males and 60% incidence in females). Similarly, sex differences in EAE severity (CDS) were noted in CC046 (greater severity in males) and CC042 (greater severity in females). These findings suggest the existence of gene-by-sex interactions, if only in a subset of the studied CC strains, the mechanisms of which can be elucidated in future studies. These findings are fully consistent with our published data in the B6.Chr^PWD^ consomic model^24^ and studies in classic inbred strains^74^, where sex differences are typically present only on a specific genetic background, e.g., in SJL and not B6 mice. As such, our studies provide new genetic models to study sex differences using a uniform EAE induction method. Given that strain-by-sex interactions were only seen in a subset of strains, our QTL mapping was performed using sex as an additive, rather than interactive covariate. Additionally, given the limited number of mice of each sex per strain, we opted not to perform sex segregated QTL analysis. However, future mapping studies could focus on subpopulations (e.g. F2 or CC-RIX) derived from CC strains exhibiting the most robust sex differences and use sex as an interactive covariate, or use sex-segregated QTL analysis, to identify specific QTL and candidate genes interacting with sex. Interestingly, a recent GWAS analysis of MS severity (rather than incidence; discussed further below), reported two sex-specific associations^41^.

One prominent advantage of the CC model is the ability to map loci controlling specific traits of interest, which we applied to different EAE phenotypes. Given the availability of extensive GWAS assessing MS incidence using case-control studies, we could leverage these results for comparison with our own EAE QTL. Candidate gene prioritization of the classic EAE incidence QTL, *Eaecc2*, identified two genes passing an FPR of 0.05, *Klf3* and *Rhoh* (**Fig. S10 C** and **D**), both of which have been associated with MS risk in the 2019 GWAS analysis^9^. Furthermore, comparison of all identified genes within EAE QTL (without prioritization) to the MS incidence GWAS, revealed additional shared genes between these analyses, including: *Txk/TXK* and *N4bp2/N4BP2* associated with *Eaecc2, Chd7/CHD7* and *Ints8/INTS8* associated with *Eaecc1* (EAE incidence), all identified in the 2019 MS GWAS^9^, as well as *Gpc5/GPC5* associated with EAE severity (*Eaecc6*), which was identified in an early MS GWAS in 2009^75^. Taken together, these results identify overlapping genes driving predominantly disease susceptibility, rather than disease course, in both EAE and MS, supporting a shared genetic architecture between the human disease and its model, in concordance with our previous EAE QTL mapping studies in SJL and B10.S mice^76,77^. With regard to our approach using machine learning-based candidate gene prioritization to identify the top positional candidate genes for each EAE QTL, we note as a caveat that the candidates may be somewhat biased towards those associated with disease risk/susceptibility rather than disease progression. The reason for this is that our approach requires a large training set of “true positive” genes (200-500 genes) known to be associated with the phenotype of interest, typically by GWAS (**Table S7**). The vast majority of these genes (available from the GWAS catalog) were associated with MS risk, although a small number also represented emerging GWAS hits for MS progression (discussed below). Therefore, our top prioritized candidate genes are likely biologically more associated with MS genes, which may be genetically distinct from progression. Hence, we consider all genes within our QTL as plausible candidates, and we can continue to improve our prioritization using larger training datasets focused specifically on progression, as those data become available in the future. While there is a wealth of genetic associations with MS risk/incidence from large case-control studies, genome-wide association studies of disease severity and/or progression have been more challenging to perform. However, new studies and associations are beginning to emerge^41,42,78,79^. We used these studies to generate a list of genes associated with MS severity/progression (See Materials and Methods; **Table S8**) and asked whether any overlapped with our EAE QTL. Of particular interest, this analysis revealed an overlapping gene from the EAE severity QTL *Eaecc6, Spry2/SPRY2,* which is the closest gene associated with a suggestive hit on Chr13 (rs2876767) from the most recent International MS genetics Consortium GWAS analysis of MS severity and progression^41^. Additionally, our comparison also revealed two overlapping genes associated with *Eaecc1* and *Eaecc2*, *Chd17/CHD17* (associated with MS severity^42^) and *Ppargc1a/PPARGC1A* (associated with accumulation of disability in MS^78^), respectively. Taken together, these studies further highlight the value of using the CC model to map genes associated with MS progression as an orthogonal approach to resource-intensive GWAS in humans. In particular, mouse QTL studies could prioritize/refine the mapping of the candidate genes from human GWAS, as in the example of *Spry2/SPRY2*, above.

Beyond overlapping MS GWAS genes, candidate gene prioritization for promising QTL, those passing 20% genome wide significance (AR-EAE incidence - *Eaecc3,* and EAE severity - *Eaecc6*) or having a particularly narrow 95% confidence interval (2.52Mb, monophasic EAE incidence - *Eaecc5*) revealed genes with relevance to MS pathology. Of note, assessment of *Eaecc6* identified *Abcc4/ABCC4,* encoding the ATP-binding cassette transporter ABCC4, as the top prioritized gene in both the immune system and CNS tissue networks. Relevant to MS pathology, ABCC4 has been implicated in blood brain barrier function^80^ and effector immune cell efflux activities, the latter of which has been shown to be involved in the pathogenesis of rheumatoid arthritis (RA)^81^, an autoimmune disease that shares a genetic burden with MS^82,83^. While the role of ABCC4 in MS is still unclear, other members of the ABCC subfamily, specifically ABCC1 and ABCC2, have been shown to involved in neuroinflammation^80^, with previous studies revealing increases in ABCC1 in the brain tissue of MS patients compared to healthy controls^84^. Taken together, these findings represent a plausible mechanism in which variants in *Abcc4/ABCC4* could contribute to increased inflammation, acting either within the CNS or peripheral immune system to drive disease severity.

In terms of AR-EAE incidence (*Eaecc3)*, *Yap1/YAP1,* which is a key component of the Hippo signaling pathway^85^, and *Dync2h1/DYNC2H1*, which encodes a dynein motor protein that has been shown to be involved in intraciliary transport, were the top prioritized genes for *Eaecc3* for the immune system and CNS networks, respectively. Recent studies have suggested the involvement of the Hippo signaling pathway in autoimmune pathogenesis, with emphasis on the balance between T regulatory cell (Treg) and pro-inflammatory Th17 cell differentiation through *Yap*-*Taz* expression^85^, as well as specific evidence for involvement in RA, inflammatory bowel disease, and psoriasis^86^, all of which have been shown to have genetic overlap with MS^82,83^. In contrast, less direct evidence for involvement in EAE/MS pathogenesis exists for *Dync2h1/DYNC2H1*. However, *Dync2h1/DYNC2H1* has been shown to have a role in neurodevelopment, maintenance, neuronal transport^87–89^, and has been shown to be associated with retinopathies^90^, suggesting a potential involvement in neurodegeneration. In further support of this, *Dync2h1* expression was downregulated in the spinal cord of mice infected with Theiler’s murine encephalomyelitis virus (TMEV)^91^, which has been used to model certain aspects of MS, including axonopathy. Taken together, these candidate genes could represent a potential mechanism behind AR-EAE incidence through immune dysregulation via various impacts of altered *Yap1* expression setting the stage for increased inflammation, and/or axonopathies via impaired transport associated with *Dync2h1* resulting in impaired neurological function.

Analysis of *Eaecc5* revealed *Il17ra/IL17RA,* encoding the receptor for the pro-inflammatory cytokines IL-17A and IL-17F, and *Wnk1/WNK1,* encoding a serine/threonine protein kinase that is involved in CNS signaling, as the top prioritized genes in the immune system and CNS networks, respectively. The association of *Il17ra/IL17RA* with monophasic EAE incidence is particularly plausible, given the known increased *Il17ra/IL17RA* expression in the CNS^92^ and involvement of IL-17 and Th17 cells in MS and EAE pathogenesis^93,94^. Specifically, studies have shown that the frequency of Th17 cells is increased during MS relapses^95^ and that assessment of serum IL-17A levels in RR-MS patients who were non-responsive to interferon-β therapy revealed that those with elevated IL-17A had shorter disease duration then those with lower IL-17A serum levels^96^. Furthermore, IL-17 has been implicated as having a significant role in the pathogenesis of both clinically isolated syndrome and early MS^97,98^. Altogether, our findings of the association of *Il17ra* with monophasic EAE incidence is in support of these previous findings, and the monophasic EAE strain CC068 represents another potential model to further elucidate the role of IL-17 in MS onset and duration. In terms of CNS expression of *Wnk1/WNK1*, it has been well accepted that *Wnk1/WNK1* is involved in pathogenic signaling in the CNS^99^ and has been implicated as a crucial component in neuronal axon development and maintenance^100^. While direct implication of *Wnk1/WNK1* in MS pathogenesis has been limited to suggestive involvement in phosphorylation differences in B cells as associated with MS susceptibility alleles^101^, the involvement of *Wnk1* in CNS dysregulation could represent a potential mechanism for association with monophasic EAE incidence. Interestingly, we identified two nonsynonymous SNPs distinguishing the two major phenotype-associated *Wnk1* alleles, both of which change the position of threonine residues (**Table 3**), providing a potential mechanism of differential post-translational regulation by upstream serine/threonine kinases^102^. Additionally, given the regulation of WNT signaling by WNK1^103^, another intriguing link between our candidate gene prioritization of *Wnk1/WNK1* and MS comes from a recent GWAS study of MS relapse hazard, which identified a minor allele in *WNT9B* as the lead candidate gene driving relapse risk^104^.

Our study is far from the first to apply forward genetics in the mouse to EAE. A number of groups have applied classic forward genetics approaches, such as F2 crosses or congenic mapping, using intercrosses between two susceptible and resistant standard inbred strains (B10.S and SJL in the majority of the studies), as reviewed in Olsson et al., 2006^105^. All together, these studies identified a large number of QTL, but, with a few exceptions^106^, the causative genes have not identified. In part, this is due to the large size of the identified EAE QTL, and thus limited resolution. The other possibility is that the polygenic nature of EAE phenotypes made it difficult to physically map candidate gene variants in cases where the genetic effects were driven by combined modest effects of several allelic variants or involved epistatic interactions^76^. Another limitation of these approaches is the use of classic inbred strains of mice, which provided limited allelic diversity, compared with CC model, which provides up to 8 different alleles, 3 of them wild-derived and divergent from classic strains. In this regard, we note that the majority of the allelic effects in our CC EAE QTL are driven by one or more wild-derived haplotypes (**Fig. 7** and **Fig. S9**), which in previous studies have been observed to exert large phenotypic effect sizes, facilitating mapping efforts^107^. The CC offers additional advantages over traditional mapping populations, including the ability to generate CC intercross (CC-RIX), or F1, mapping populations, and using in parallel the Diversity Outbred (DO) population, which carries the same founder allelic variants but with a very high degree of recombination, allowing for higher resolution mapping comparable to human GWAS^27^. This, combined with the growing availability of bioinformatic tools and shared resources, and extensive and still growing human GWAS databases, makes the CC and the DO promising systems genetics tools to continue to dissect the genetic architecture of MS and other complex autoimmune diseases, as evidenced by a recent study of systemic autoimmunity induced by silica exposure in the DO^108^.

Our study is the first (to our knowledge) to apply the CC resource in an autoimmune model of MS. However, the Threadgill and Brinkmeyer-Langford groups have utilized the CC to study neurological disease induced by TMEV CNS infection in a number of studies^109–113^. These investigators included strains of multiple *H2* haplotypes, and thus the overlap with strains in our own study is limited. However, they did identify CC002 and CC011 as strains with minimal neurological progression^109^, which is somewhat in line with the EAE phenotypes of these strains (RR-EAE and resistant, respectively). Additionally, sex differences in TMEV induced disease also varied by strain^109^, in line with our findings in EAE. While these studies were likely underpowered for QTL mapping (ranging from 5-18 CC and/or CC-RIX strains per study), and complicated by the fact that host genetic variation impacts viral clearance in this model, they nonetheless provide strong evidence that neurological sequelae following adaptive immune inflammation and demyelination in the CNS are genetically regulated, in full agreement with our own EAE studies.

Our identification and initial characterization of EAE phenotypes in CC mice represents what we hope will be the first of many utilizing the CC resource in the pursuit of the genetic underpinnings behind MS disease heterogeneity. An obvious extension of this will include the validation and functional characterization of the identified candidate genes, as well as further mapping efforts to increase the resolution and statistical significance of EAE QTL. Because expression QTL (eQTL) often underlie phenotypic QTL, and as such are often used to help fine-mapping in human GWAS, future EAE QTL mapping could be facilitated by the generation of immune and CNS cell specific eQTL databases for CC and Diversity Outbred populations. Similarly, PheWAS approaches and/or strain correlations could be used to find colocalized traits and intermediate phenotypes, such as immune or neurological phenotypes, with the latter enhanced by the ever growing immune phenotyping studies in CC strains as a key resource^36^. Other efforts will focus on further mechanistic characterization of specific EAE disease course attributes unique to subsets of CC strains as described here. Importantly, the work presented here provides to the research community an easily accessible improved animal model of MS that accounts for host genetic diversity and captures novel phenotypes lacking in traditional models.

## Materials and Methods

### Mice

For peptide-based EAE studies, male and female mice of each of the 32 Collaborative Cross (CC) strains (**Table 1**) were purchased between 2021 and 2022 from the Mutant Mouse Resource and Research Center (MMRRC) at University of North Carolina at Chapel Hill, an NIH-funded strain repository, in collaboration with the UNC Systems Genetics Core Facility (UNC; Chapel Hill, North Carolina, USA), with the exception of CC020, which was purchased from The Jackson Laboratory (Bar Harbor, Maine, USA). While CC strains were originally generated and bred at Oak Ridge National Laboratory (USA), the International Livestock Research Institute (Kenya)/Tel Aviv University (Israel), or Geniad Ltd (Australia),all CC strains had been maintained at UNC (or transferred from UNC to The Jackson Laboratory and then back to UNC for clean rederivation) for 10 years from these original diverse locations^114–117^. All CC strains are referred to in the text by their abbreviated name (“CC###”); full strain names are provided in Table 1. Due to availability and the number of mice studied, mice were obtained and studied across four cohorts – denoted as C1 – 4 (**Table 1**). To reduce confounding by batch effects, male and female B6 mice were purchased from Jackson Laboratory to serve as reference controls in each cohort. Once at the vivarium at the University of Vermont, mice were rested for a range of 14 – 29 days, depending on quarantine requirements, prior to experimentation. For follow up studies utilizing CC002, CC004 and CC028, including CNS flow cytometry and reciprocal bone marrow chimeras, three females and two males of each strain were purchased from the MMRRC in collaboration with the UNC Systems Genetics Core Facility in 2022 and bred at the vivarium at the University of Vermont until sufficient numbers of offspring were obtained for experimentation. The experimental procedures used in this study were approved by the Institutional Animal Care and Use Committee of the University of Vermont.

### Selection of CC Strains for Peptide-induced EAE

To identify CC strains that are compatible with myelin oligodendrocyte glycoprotein 35-55 peptide (MOG_35-55_) induced experimental autoimmune encephalomyelitis (EAE), the *H2* haplotype of each strain was characterized. Founder strain contributions at the H2 locus, specifically in the region encompassing MHC class I (*H2^K^* only) and all class II genes, as defined by Chr17:33,918,830–34,347,345 bp, were identified using the UNC Systems Genetics Collaborative Cross Viewer tool (Chapel Hill, North Carolina, USA), using sequenced and most recent common ancestor (MRCA) genotypes. Given that previous studies have suggested MOG_35-55_ binding compatibility with *H2^b^* (C567BL/6J (B6) and 129S1/SvlmJ (129S1)) and *H2^g^*^7^ (NOD/ShiLtJ (NOD)) haplotypes^44–46^, CC strains with founder contributions, either homozygous or heterozygous, from the three abovementioned strains were selected for study, for a total of 32 investigated CC strains (see **Table 1** for strain list and corresponding *H2* haplotype).

### Mouse Spinal Cord Homogenate Induced EAE Studies

To investigate the use of mouse spinal cord homogenate (mSCH) as a potential EAE immunogen in CC mice, three CC strains (CC003/Unc, CC008/GeniUnc, and CC010/GeniUnc) with different *H2* haplotypes, and four CC founder strains (129S1, A/J, B6, and NOD) were selected for analysis. CC mice were purchased in 2020 from the UNC Systems Genetics Core Facility^114,116^, then bred and housed within the vivarium at the University of Vermont until sufficient numbers were obtained for experimentation. CC founder strains were purchased from The Jackson Laboratory and then rested for two weeks prior to experimentation.

mSCH was prepared from Swiss Webster mice (Charles River Laboratories) and EAE was induced using a modified approach from previously described^22,25^. Briefly, mice were injected subcutaneously with 0.15 ml of an emulsion containing 5 mg mSCH in PBS and 50% complete Freund’s adjuvant (CFA; Sigma, USA), supplemented with additional 4 mg/ml heat-killed *Mycobacterium tuberculosis* (Difco Laboratories, USA). On D0 or D0 and D2, mice were administered an intraperitoneal injection of 200 ng pertussis toxin (PTX; List Biological Laboratories, USA) as an ancillary adjuvant. At 7 days post induction, mice were scored daily for presence of clinical disease symptoms as previously described^22,24^, and expanded below.

### Peptide-based EAE Induction and Scoring

Approximately five male and five female mice, 8-14 weeks old, of each of the 32 CC strains were immunized subcutaneously with 200 µg MOG_35-55_ (New England Peptide) emulsified in CFA (Sigma), supplemented with additional 4 mg/ml heat-killed *Mycobacterium tuberculosis* (Difco Laboratories), and received a single intraperitoneal injection of 200 ng PTX (List Biological Laboratories) on D0 as an ancillary adjuvant. Mice were studied across a total of four cohorts. Five male and five female B6 mice per cohort, immunized as above, served as reference controls for cohort effects and EAE induction.

Mice were observed daily for a total of 50 days starting at 5 days post induction for the presence of clinical disease symptoms, using both classic EAE^22,24^ or a modified axial rotary (AR)-EAE scoring scale^43^. Briefly, classic EAE clinical scores were assigned as follows: 0, asymptomatic; 1, tail paralysis; 2, tail paralysis and hind limb weakness; 3, hind limb paralysis; 4, hind limb paralysis with incontinence; and 5, moribund/quadriplegic. AR-EAE clinical scores were assigned as follows: 0, asymptomatic; 1, slight head tilt; 2, pronounced head tilt; 3, inability to walk in a straight line; 4 mouse is moving/lying on its side, will continuously fall to its side after being made to stand; 5, mouse rolls or spins continuously - axial rotation. Mice reached humane endpoints after presenting with a score of 5 (classic or AR-EAE) for 72 hours, at which point mice were euthanized and their daily disease score was recorded as a 5 (classic or AR-EAE depending on presentation) for the remainder of the experiment. Similarly, mice that died after having presented with EAE for more than two continuous days were given a daily disease score of 5 for the remainder of the experiment (designated as classic or AR-EAE depending on previous presentation). Mice that died without any EAE clinical signs or having only presented with EAE for two or fewer days were excluded from the study.

### EAE Disease Phenotype/Quantitative Trait Variable (QTV) Classification

Raw daily disease scores, reported as both classic and AR-EAE as described above, for the full 50-day time course were pooled by strain and sex upon completion of all four cohorts. Raw daily disease scores were utilized to derive the following daily score classifications: classic disease score (reports disease scores that only correspond to the classic-EAE scoring system and negates/assigns a score of 0 to any AR score if present in the raw score; **Table S3**), AR disease score (reports disease scores that only correspond to the AR-EAE scoring system and negates/assigns a score of 0 to any classic score if present in the raw score; **Table S4**), and disease score (reports disease scores regardless of EAE subtype and derives an average score for any occurrence of simultaneous classic and AR-EAE scoring, i.e. a mouse that was reported to have a classic score of 3 and an AR score of 2 on the same day was assigned a score of 2.5; **Table S2**). Subsequently, these three disease score classifications (disease score, classic disease score, and AR disease score) were utilized to determine cumulative disease score (CDS; the total sum of all daily disease scores), classic cumulative disease score (classic-CDS; the total sum of all daily classic disease scores), and AR cumulative disease score (AR-CDS; the total sum of all daily AR disease scores), respectively. CDS, classic-CDS, and AR-CDS were calculated per mouse and averaged by sex and strain to allow for analysis of both strain and sex effects.

To assess incidence of EAE and EAE subtypes, the following quantitative trait variables (QTVs) were calculated per individual mouse: EAE incidence, classic EAE incidence, AR-EAE incidence. EAE incidence was determined utilizing daily disease scores and classified as a minimum of two consecutive days of a score above 0. Classic EAE incidence and AR-EAE incidence were assessed in a binary and mutually exclusive manner, as determined by raw daily disease score. First, mice were categorized into three groups: classic EAE only (mice that only presented with clinical presentations aligning with the classic EAE scoring system), AR-EAE only (mice that only presented with clinical presentations aligning with the AR-EAE scoring system), and “combined type” (mice that presented with clinical presentations of both the classic and AR-EAE scoring system, either on the same or different days during the experiment). Mice that were classified as being in the classic EAE only or AR-EAE only group were assigned as either classic EAE incidence or AR EAE incidence, respectively. For mice classified as being in the “combined type group”, EAE subtype incidence was assigned as classic EAE incidence if the number of days in which a mouse presented with only a classic EAE score (excluding days of simultaneous classic and AR-EAE scores) was greater than the number of days in which a mouse presented with only an AR-EAE score (excluding days of simultaneous classic and AR-EAE scores). The inverse was utilized to assign AR-EAE incidence. For all incidence calculations, individual mice were assigned a value of 0 or 1 given the absence or presence of the assigned trait, respectively. To assess incidence by strain and sex, values (assigned 0 and 1) per QTV were tallied by either strain or sex and then divided by the corresponding number of mice per strain or sex and multiplied by 100 to calculate percent incidence.

To determine EAE disease course phenotypes, individual mice were classified as having either a relapsing remitting (RR), monophasic, or chronic EAE disease course as assessed using daily disease scores (**Table S2**). RR-EAE was defined as a scoring pattern in which a mouse presents with an initial disease period (defined by at least two consecutive days of a disease score >0), followed by remission period (characterized by disease score of 0 for three or more consecutive days), and proceeded by a relapse period of at least two consecutive days of a disease score >0. Mice characterized as having RR-EAE could have one or more remission and relapse cycles. Classification of monophasic EAE was defined as a scoring pattern in which a mouse presented with a consistent disease score of 0 after having presented with a disease score >0 for at least two consecutive days. Chronic EAE was defined as a scoring pattern in which a mouse presented with persistent disease scores >0 from time of disease onset until experiment termination (or humane endpoint) that could not be classified as either RR-EAE or monophasic EAE. Percent incidence of RR, monophasic, and chronic EAE by strain and sex was determined as described above.

### CNS Histopathology

On day 50 post EAE induction, or upon reaching humane endpoint criteria, mice were euthanized, and brain and spinal cord tissues were collected for histopathological assessment as previously described^49^. In brief, the skull and vertebral column were removed, and diffusion fixed in 10% neutral buffered formalin (Fisher Scientific, USA). Post fixation, brain and spinal cord were extracted from calvaria and vertebral columns, respectively, and dissected into thirds based on overarching anatomical region (brain: front brain, mid brain, and hind brain – including cerebellum; spinal cord: cervical, thoracic, and lumbar). Tissues were subsequently embedded in paraffin, sectioned (coronal and longitudinal for brain and spinal cord, respectively), and stained with luxol fast blue (LFB) and/or hematoxylin and eosin (H&E) by the pathology department at the University of Vermont Medical Center.

Histopathological assessment was conducted in a blinded fashion and evaluated representative areas of the brain and spinal cord corresponding to the above-mentioned regions. Brain and spinal cord regions were assessed for degree of inflammation and demyelination using a semi-quantitative scale adapted from previous studies^49,50^. Inflammation was evaluated using H&E stained tissues and scored on a scale of 0 – 4 based on severity using the following criteria: 0, no inflammation; 1, few inflammatory cells scattered or in small clusters; 2, organized clusters of inflammatory cells without significant extension beyond small lesions; 3, significant organized clusters of inflammatory cells with patchy infiltration of surrounding tissue, central involvement of larger lesions; and 4, extensive and dense infiltration of inflammatory cells with affecting over half of the sample, +/− diffuse gliosis. Extent of demyelination was evaluated from tissues stained with LFB (co-stained with H&E) and scored on a scale of 0 – 4 based on severity using the following criteria: 0, no demyelination – deep blue staining; 1, small, patchy area(s) of white matter pallor on LFB, no well-defined lesions; 2, defined area of white matter pallor forming isolated lesion(s); 3, confluent foci of white matter pallor on LFB with some spared areas in the sample; and 4, widespread white matter pallor on LFB affecting nearly all of the sample (over ∼75%). Inflammation and demyelination scores were assigned to each of the three dissected regions per tissue (brain and spinal cord)/mouse. Overall inflammation and demyelination scores were reported per mouse for both brain and spinal cord and were determined by the highest scored region of that tissue.

### Flow Cytometry

To assess CNS infiltrating cells, mice were anesthetized under isoflurane, euthanized by exsanguination via transcardial perfusion with PBS, and brain and spinal cord tissues were collected and processed independently for flow cytometric staining as previously described^118,119^. In brief, brain and spinal cord tissues were mechanically dissociated by Dounce homogenization until a single cell suspension was created. Resulting cell suspensions were filtered with a 70-μm strainer followed by Percoll gradient (37%/70%) centrifugation and collection of the interphase for subsequent processing dependent on the objective of flow cytometry analysis.

For intracellular cytokine analysis, cells were stimulated with 5 ng/mL PMA, 250 ng/mL ionomycin, and brefeldin A (Golgi Plug reagent; BD Bioscience, USA) for 4 hours prior to staining. For flow cytometry analysis of intracellular cytokines and surface staining, cells were stained with UV-Blue LIVE/DEAD fixable stain (Invitrogen, USA) and then surface stained with antibodies against the following: CD45, CD19, CD11b, TCRβ, CD4, CD8, Ly6G, and CX3CR1 (BioLegend, USA). Cells were then fixed and permeabilized with 1% paraformaldehyde (Sigma Aldrich, USA) and 0.05% saponin permeabilization buffer and labeled with antibodies against IFNγ, IL-17A, and GM-CSF (BioLegend).

For flow cytometry staining to assess chimerism in addition to CNS infiltrating cells in CC002 and B6 mice, spleens were collected at time of euthanasia in addition to spinal cord. Spinal cord tissues were processed as described above and spleens were processed as previously described^118^. Briefly, spleens were mechanically dissociated by a syringe plunger between two pieces of mesh to create a single cell suspension followed by red blood cell lysis by incubation in 0.8% ammonium chloride solution (STEMCELL Technologies) and subsequent staining. Cells were stained with UV-Blue LIVE/DEAD fixable stain (Invitrogen) and then surface stained with antibodies against the following: CD45.1, CD45.2, CD19, CD11b, TCRβ, CD4, CD8, Ly6G, and CX3CR1 (BioLegend, USA) prior to fixation with 1% paraformaldehyde (Sigma Aldrich).

All stained cells were analyzed utilizing a Cytek Aurora and SpectroFlo software versions 2.2 – 3.3 (Cytek Biosciences, USA) and spectral unmixing was performed with appropriate single-color controls and autofluorescence correction from an unstained control group. Flow cytometry data analysis was performed using FlowJo software versions 10.8.1 – 10.10 (BD Biosciences).

### Reciprocal Bone Marrow Chimeras

To identify CD45 allele status in CC002 mice, founder strain contributions for the gene *Ptprc* (Chr1:137,990,599-138,103,446 bp) were identified using the UNC Systems Genetics Collaborative Cross Viewer tool (Chapel Hill, North Carolina, USA), using both sequenced and MRCA genotypes. Recipricoal bone marrow chimeras were generated between B6.SJL-Ptprca Pepcb/BoyJ (B6.CD45.1) and CC002 mice as previously described^118^. In brief, 9-15 week old recipient mice were irradiated twice with 550 rads 4-6 hours apart. Post irradiation, mice were injected via retro-orbital vein with 4×10^6^ whole bone marrow cells from respective unmanipulated age- and sex-matched donors. Resulting chimeric mice were rested for 8 weeks to allow for maximal immune reconstitution, at which point EAE was induced as described above and disease course was observed for a total of 30 days. At 30 days post EAE induction, mice were euthanized, and spleen and spinal cord tissues were collected to assess rate of chimerism and CNS immune cell profiles via flow cytometry as described above.

### Quantitative Trait Loci (QTL) Mapping

To map genetic variants that are associated with EAE QTV (including: CDS, total EAE incidence, AR-EAE incidence, classic EAE incidence, RR-EAE incidence, monophasic EAE incidence, and chronic EAE incidence), the R package, R/qtl2^52^, was utilized to preform quantitative trait loci (QTL) mapping. QTVs were calculated as described above. For QTL mapping with CDS, individual mouse data was utilized and underwent covariate batch correction, using experimental batch as a covariate, and rank Z normalization to address potential batch effects. For QTL mapping addressing incidence QTVs, strain incidence percentages were utilized, and rank Z normalization was performed. CC genotype probabilities and kinship matrices were derived utilizing the CC genome “sequenced” data set available from the UNC Systems Genetics Core Facility (Chapel Hill, North Carolina, USA (http://csbio.unc.edu/CCstatus/CCGenomes/%23genotypes). Resulting Manhattan plots demonstrating logarithmic of odds (LOD) traces and significance thresholds of 20% genome wide significance were generated given analysis utilizing 1000 permutations. Allele effect plots were generated for lead QTL for each QTV. For analysis of large QTL effects, QTV specific allele effect plots were generated in addition to classic allele effect plots, allowing for subsequent CC strain-specific genotype by phenotype analysis.

### Candidate Gene Prioritization

To identify and prioritize potential causative genes associated with the *Eaecc* QTL (**Table 2**) in an unbiased manner, we leveraged a machine learning-based approach that we developed and described previously^55–57^. We trained support vector machines (SVM) to classify randomly selected genes from previously identified MS GWAS genes as reported by the National Human Genome Research Institute GWAS catalog^59^. We used the top 500 as determined by -log10(p value) for training (**Table S7**). Mouse orthologs were identified for all genes in the training set and positional candidates were removed so as not to be used for SVM training. Feature vectors for training the SVMs were based on connection weights in one of two functional networks of tissues in mouse (FNTM) network – hemolymphoid system, as a proxy for the immune system, or CNS. The feature vector for a single true positive gene consisted of its connection weights to all other true positive genes in the training set. Any genes without connections to other genes in training set were trimmed off, resulting in a total of 271 and 273 positive-labeled genes for training the immune system and CNS networks, respectively (**Table S7**). For each tissue-specific network we trained 100 independent SVMs. Each SVM was trained to classify the roughly 271 positively labeled MS genes from a matched set of randomly drawn genes from outside the set of GWAS candidates. All trained SVMs were then used to classify positional candidate genes in each QTL. The final score for each gene was the -log_10_ of it false positive rate (FPR) averaged across all 100 SVMs. FPR for gene *x* was defined as the following: FPR_x_ = FP/(FP + TN), where FP is the number of false positive genes and TN is the number of true negative genes using a cutoff score equal to that of gene *x*. Significance for prioritization analysis was determined using an FPR of 0.05.

To determine direct overlap between the identified candidate genes associated with EAE QTL and mouse orthologs of genes associated with MS risk by GWAS, we compared all genes within a given EAE QTL to the complete list of reported genes from the 2019 GWAS analysis^9^, as well as the genes associated with MS susceptibility from the original list of the top 500 genes used in the SVM training pipeline (**Table S7**). Similarly, we generated a list of mouse orthologs of genes associated with MS severity/progression based on recent studies^41,42,78,79^ (**Table S8**), to determine overlap between the identified candidate genes associated with EAE QTL and genes associated with MS severity.

### Statistical Analysis

Statistical analysis not pertaining to QTL mapping and candidate gene prioritization were carried out using GraphPad Prism software, versions 9.1.2 – 10.2.1. Details of the analyses are provided in the figure legends; including the specific tests used to assess the significance of the observed differences and indication of adjustments for multiple comparisons when appropriate. Comparisons were assessed for effects between B6 and CC strains or within strain sex effects as indicated. All center values represent the mean, and error bars represent the standard error of the mean. A P value <0.05 was considered significant. Comparisons are indicated by the brackets and P values are reported using asterisks where significant (* p ≤ 0.05, ** p ≤ 0.01, *** p ≤ 0.001, **** p ≤ 0.0001). A lack of an indicated asterisk p value therefore signifies a lack of a significant difference.

## Data Availability

The data underlying the figures and tables presented are available in this article and its supplementary material.

## Acknowledgements

The authors would like to acknowledge the University of Vermont Medical Center Histology Lab, the Flow Cytometry and Cell Sorting facility, the Department of Pathology and Laboratory Medicine at the University of Vermont Larner College of Medicine for the use of their facilities and resources. They also acknowledge the University of North Carolina Systems Genetics Core Facility, and in particular the core director Darla Miller for her advice and help coordinating mouse experiments. Additionally, the authors would also like to specifically acknowledge Dr. DeWitt in the Department of Pathology and Laboratory Medicine for consulting on the guidelines used for histopathology scoring, as well as the members of the Krementsov Lab over the course of this study (Theresa Montgomery, Ph.D, Bristy Sabikunnahar, Ph.D, Dan Peipert, and Sydney Caldwell) for their assistance in tissue harvests.

This work was supported by RG-1901-33309 from the National Multiple Sclerosis Society as well as R21AI145306 and R01AI172166 from the NIH | National Institute of Allergy and Infectious Diseases to DNK. With additional support by U19AI100625 from the NIH | National Institute of Allergy and Infectious Diseases to MTF. Additional support for EAH was provided by 5T32AI055402-08 from the NIH | National Institute of Allergy and Infectious Diseases. The distribution of the CC mice used in this study was supported by U42OD010924 from the NIH to the Mutant Mouse Resource and Research Centers (MMRRC) at UNC.

## Author Contributions

DNK, EAH, JMM, MTF, RML, and CT designed the research; EAH, AT, TL-W, KGL, and KH performed the research; EAH, AT, MTF, JMM, and DNK analyzed the data and interpreted results; EAH and DNK wrote the manuscript; EAH, DNK, AT, TL-W, MTF, RML, and CT edited the final manuscript.

