## Supplemental Figures for "Probing the basis of disease heterogeneity in multiple sclerosis using genetically diverse mice"

### Figure S1

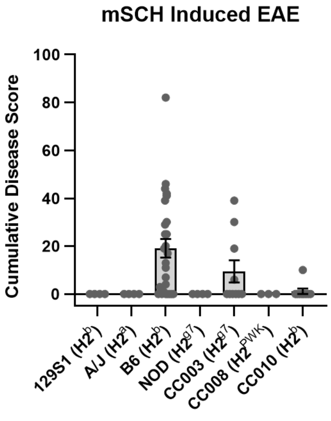


**Figure S1. mSCH-based induction of EAE in CC mice with various *H2* haplotypes is inefficient.** EAE was induced 8-20 week old mice with various H2 haplotypes including: CC founder strains - 129S1 ($H2^b$; 4M), A/J ($H2^a$; 4M), B6 ($H2^b$; 17M + 10F), and NOD ($H2^{g7}$; 4M), and CC strains: CC003 ($H2^{g7}$;8M + 2F), CC008 ($H2^{PWK}$; 3F), and CC010 ($H2^b$; 2M + 7F) by s.c. immunization with 0.15 ml of an emulsion containing 5 mg mSCH in PBS and 50% CFA. On D0 or D0 and D2, mice were administered an i.p. injection of 200 ng PTX as an ancillary adjuvant (see Materials and Methods). Mice were observed daily for a total of 26 days starting at 7 days post induction for the presence of clinical disease symptoms. (**A**) Cumulative disease score of above-mentioned strains shown with bars to demonstrate strain averages and points to display individual mice.

### Figure S2 1/2

#### Disease Course – Strain Average

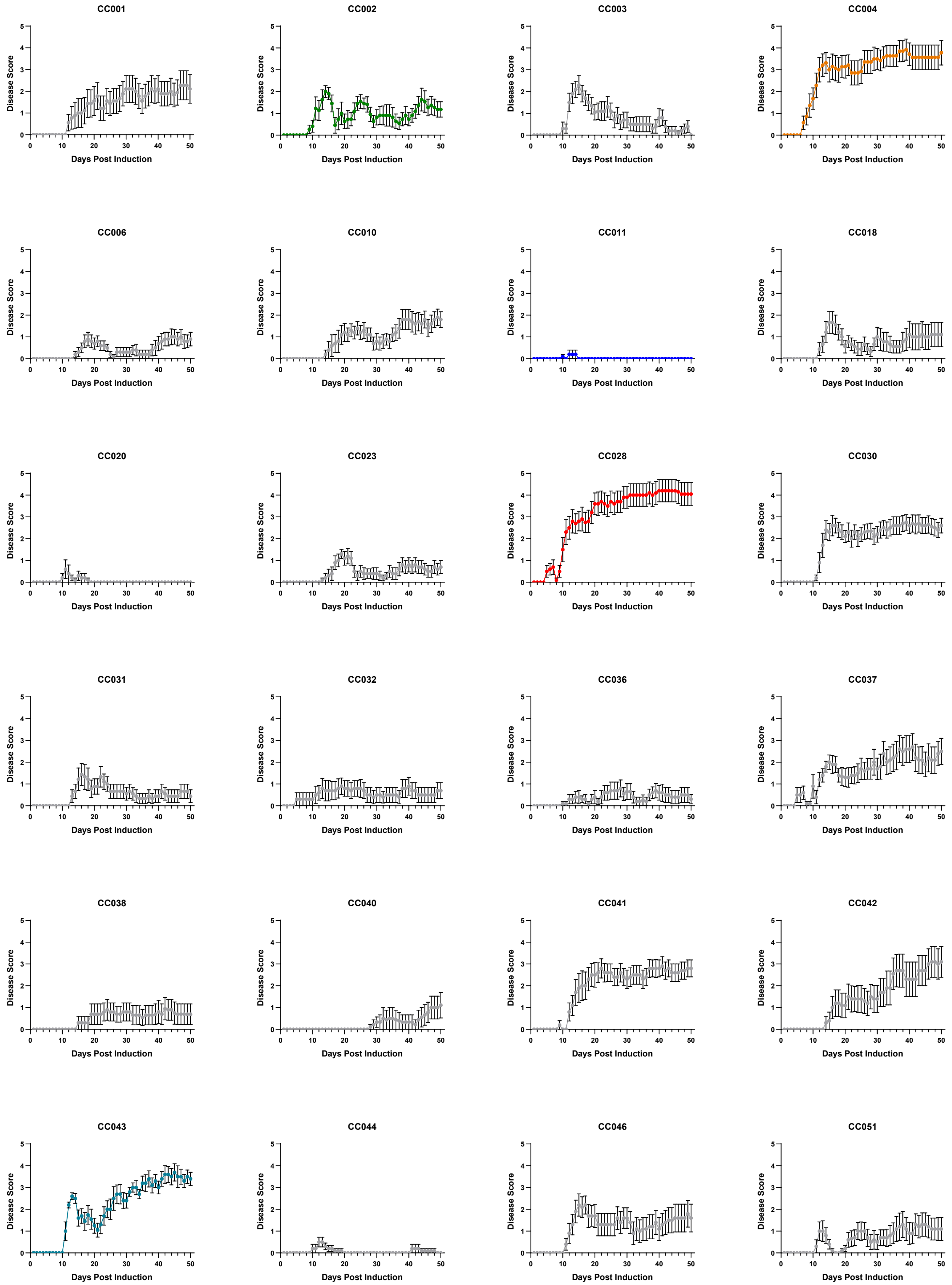

### Figure S2 2/2

#### Disease Course – Strain Average

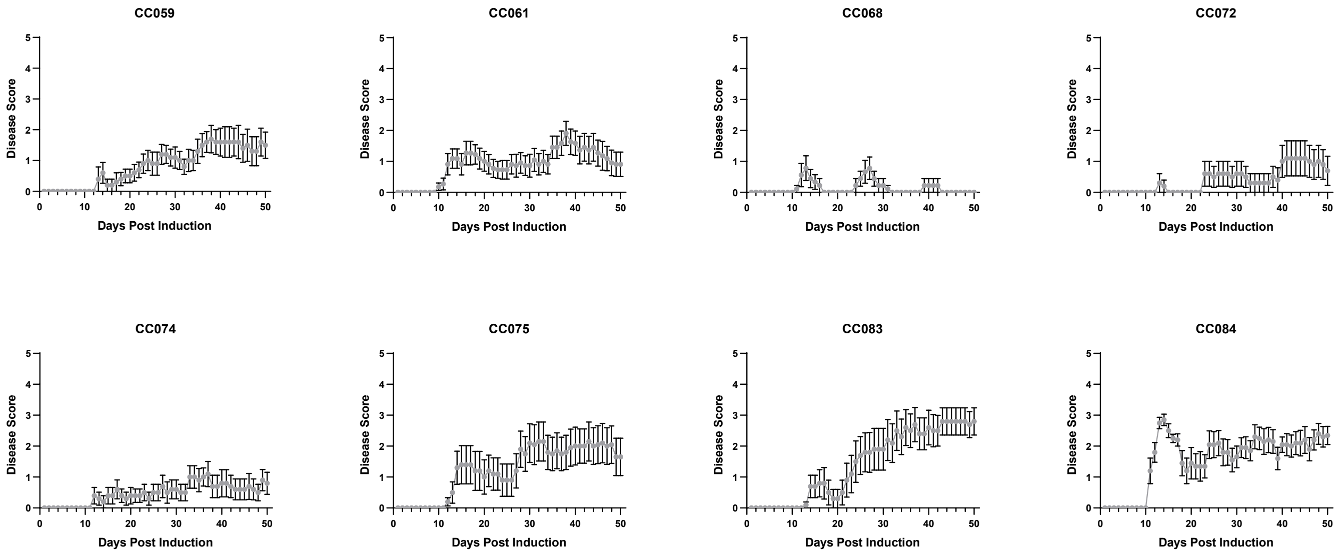


**Figure S2. Disease course profiles for CC strains.** EAE was induced and observed for 50 days in CC mice as described in Figure 1. Post observation, disease course profiles for each strain were derived from daily disease scores (see Materials and Methods). Disease course profiles as calculated by strain average for each CC strain are displayed. CC strains are displayed in numerical order by strain number and strains highlighted in Figure 2 retained strain specific label coloring.

### Figure S3 1/2

#### Disease Course – Sexes

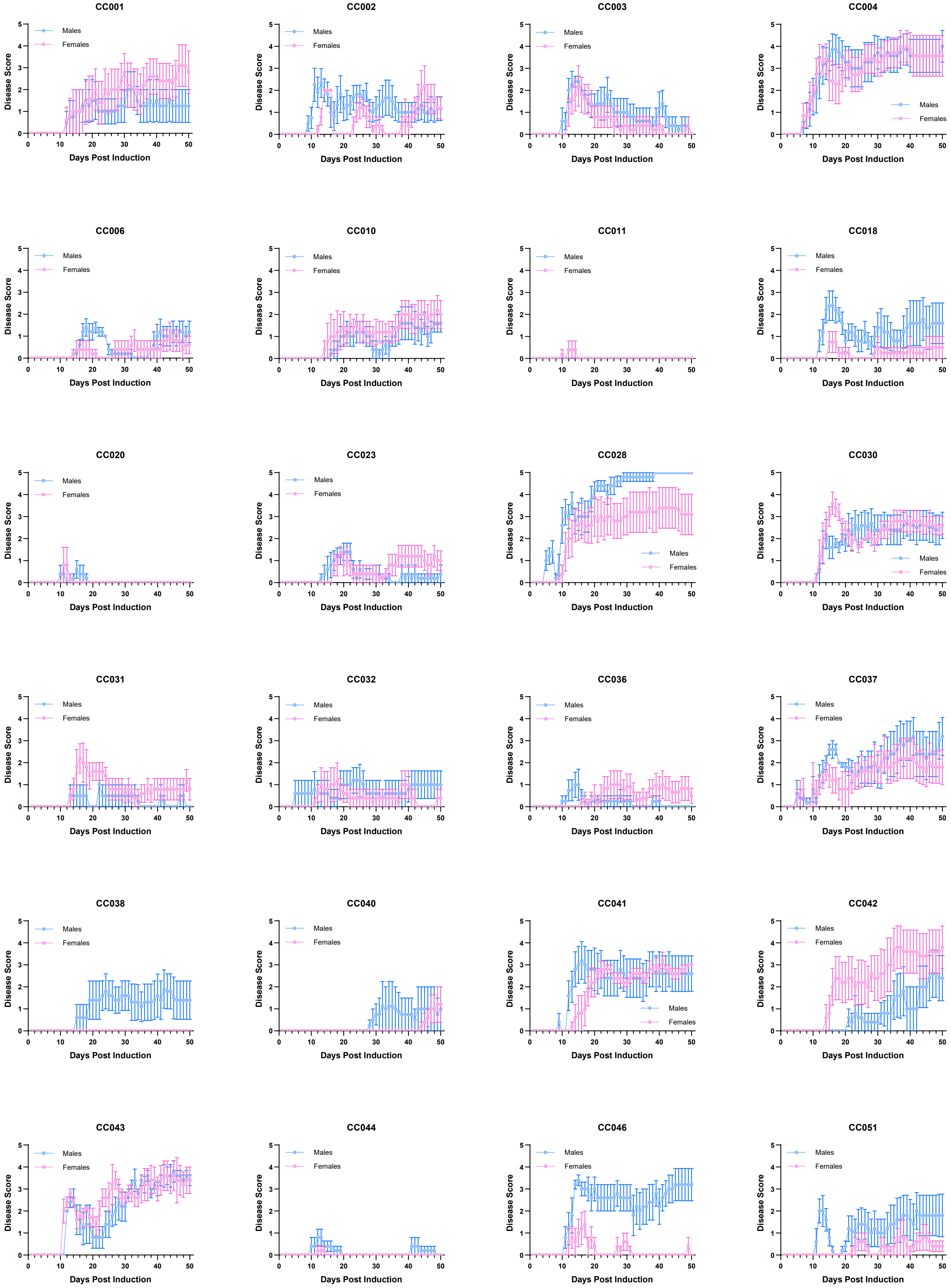

### Figure S3 2/2

#### Disease Course – Sexes

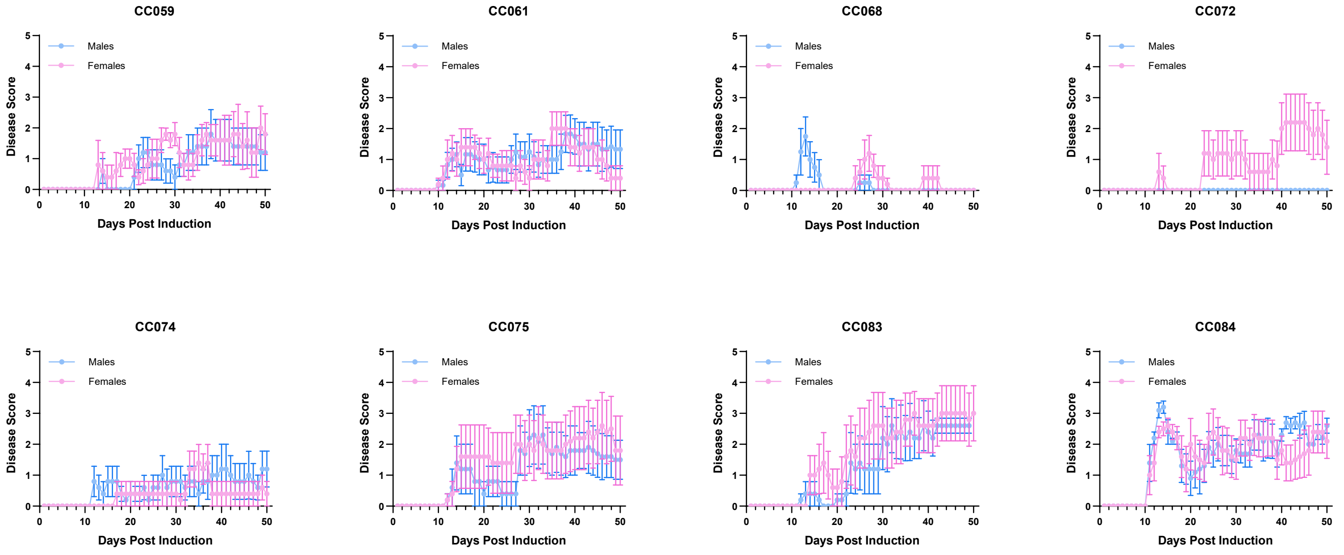


**Figure S3. Disease course profiles for CC strains separated by sex.** EAE was induced and observed for 50 days in CC mice as described in Figure 1. Post observation, disease course profiles for each strain were derived from daily disease scores (see Materials and Methods). Sex specific disease course profiles as calculated by male (blue) and female (pink) averages for each CC strain are displayed. CC strains are displayed in numerical order by strain number.

### Figure S4 1/2

#### Classic Disease Course – Strain Average

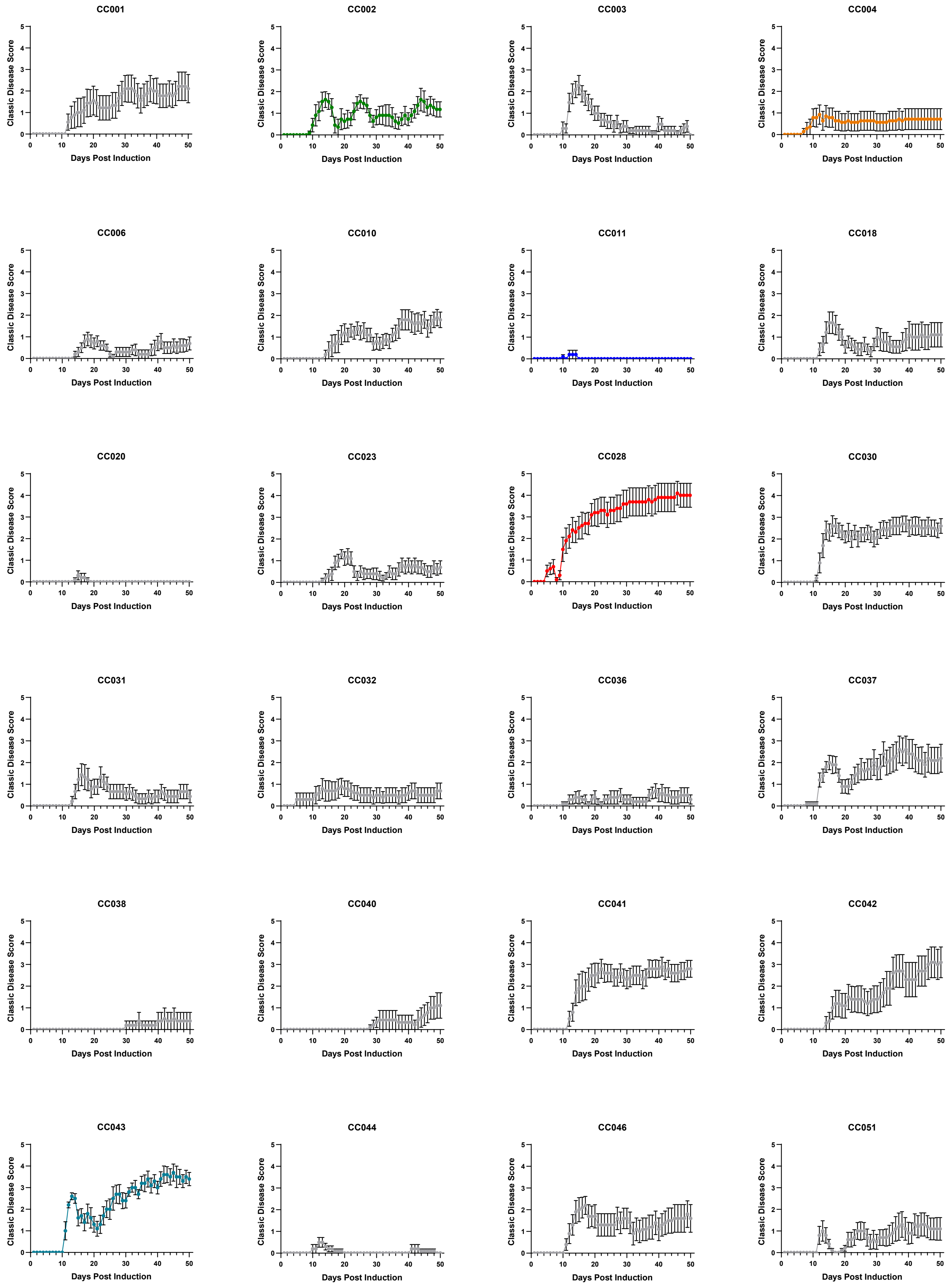

### Figure S4 2/2

#### Classic Disease Course – Strain Average

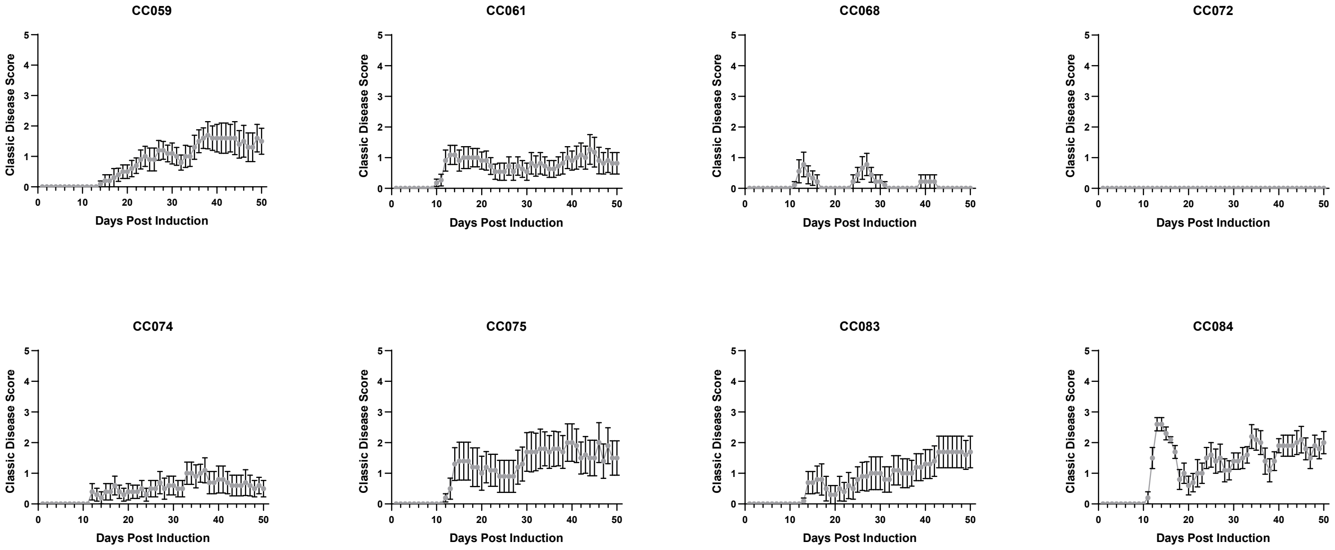


**Figure S4. Classic EAE disease course profiles for CC strains.** EAE was induced and observed for 50 days in CC mice as described in Figure 1. Post observation, disease course profiles for each strain were derived from daily classic EAE disease scores (see Materials and Methods). Classic EAE disease course profiles as calculated by strain average for each CC strain are displayed. CC strains are displayed in numerical order by strain number and strains highlighted in Figure 2 retained strain specific label coloring.

### Figure S5 1/2

#### Classic Disease Course – Sexes

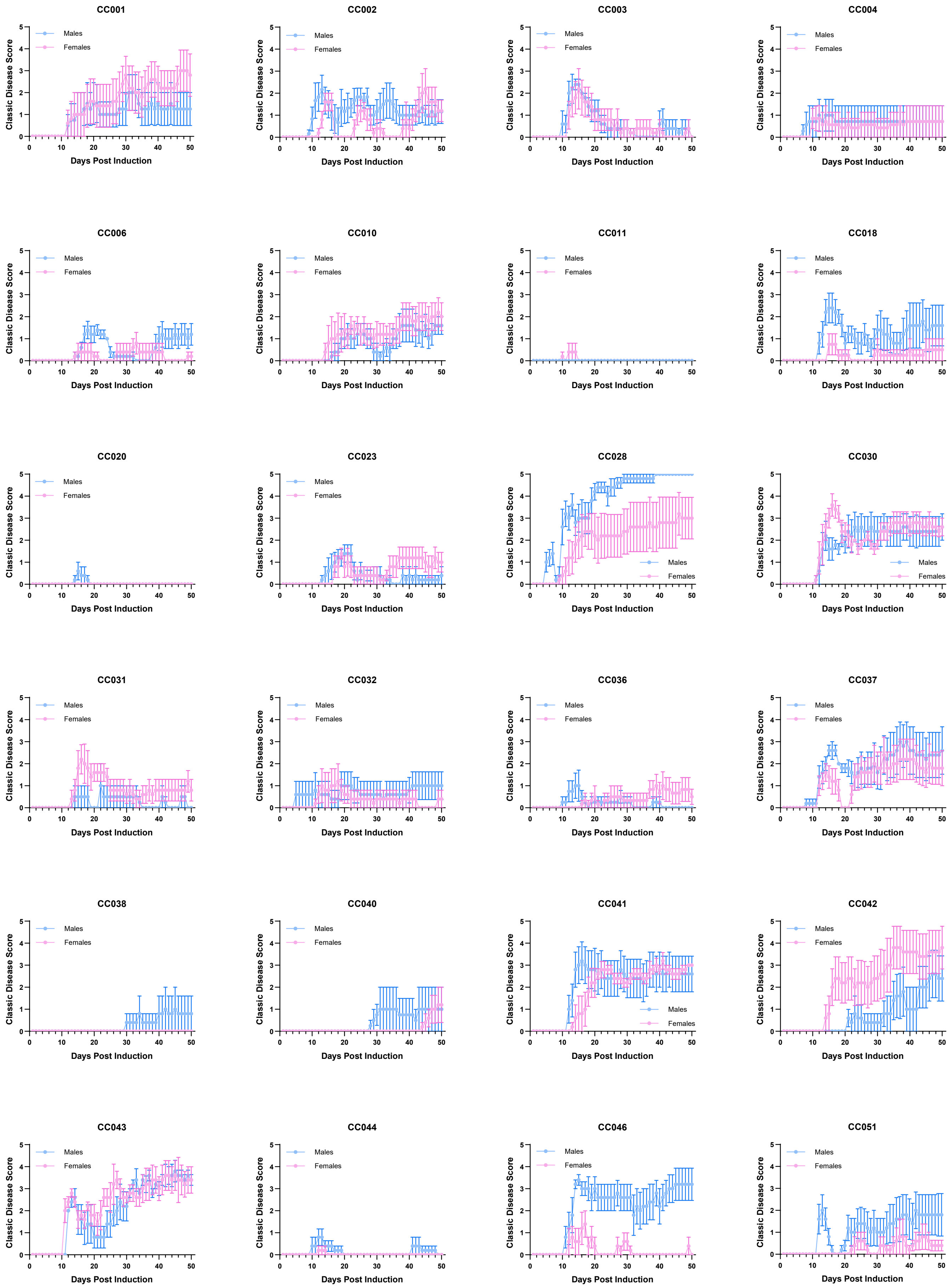

### Figure S5 2/2

#### Classic Disease Course – Sexes

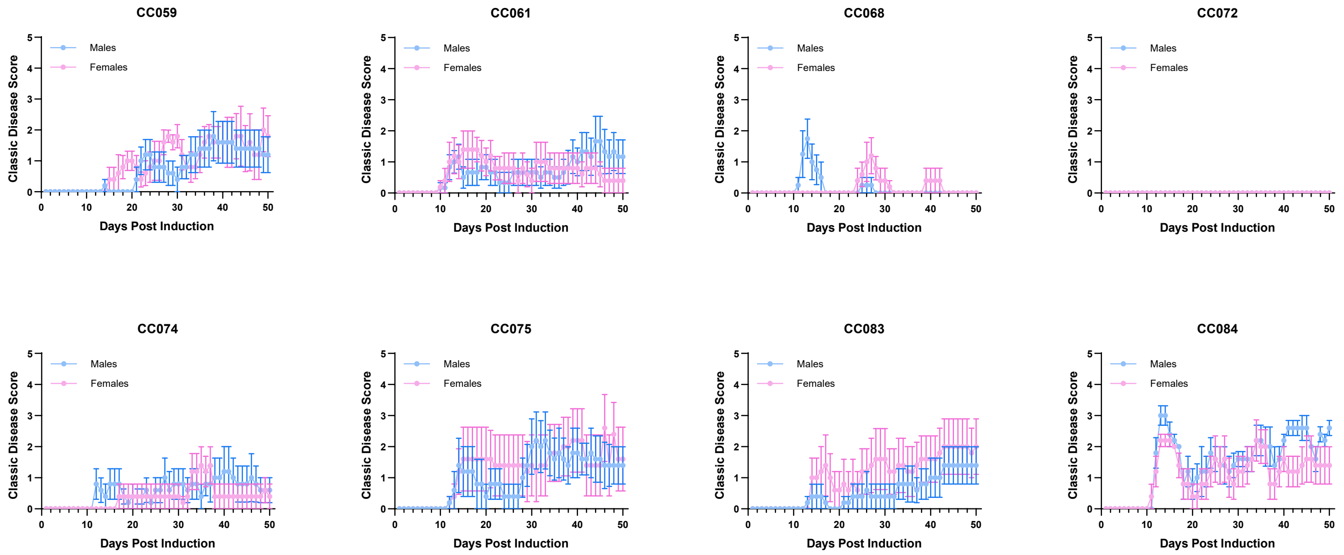


**Figure S5. Classic EAE disease course profiles for CC strains separated by sex.** EAE was induced and observed for 50 days in CC mice as described in Figure 1. Post observation, disease course profiles for each strain were derived from daily classic EAE disease scores (see Materials and Methods). Sex specific classic EAE disease course profiles as calculated by male (blue) and female (pink) averages for each CC strain are displayed. CC strains are displayed in numerical order by strain number.

### Figure S6 1/2

#### Axial Rotary Disease Course – Strain Average

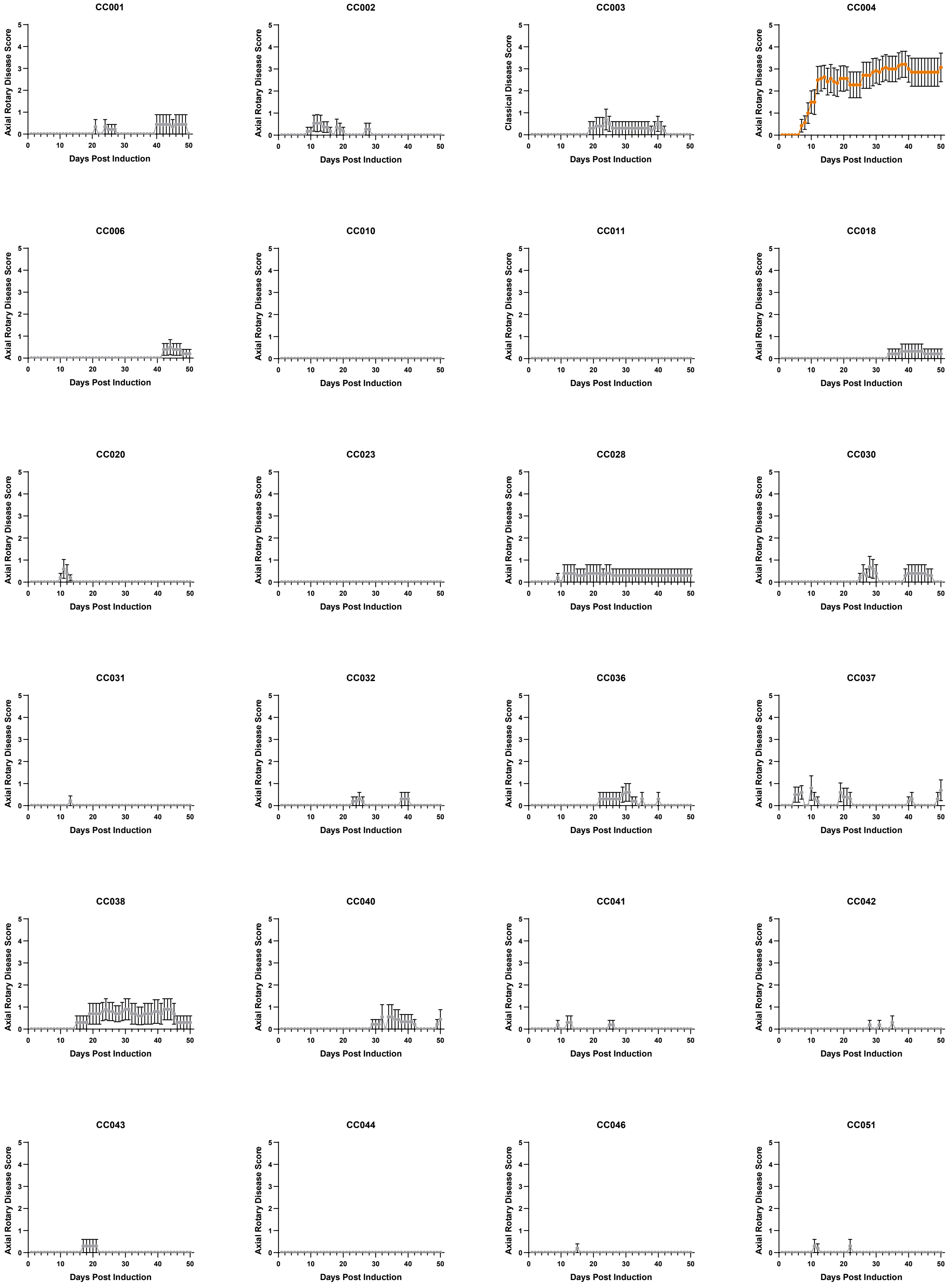

### Figure S6 2/2

#### Axial Rotary Disease Course – Strain Average

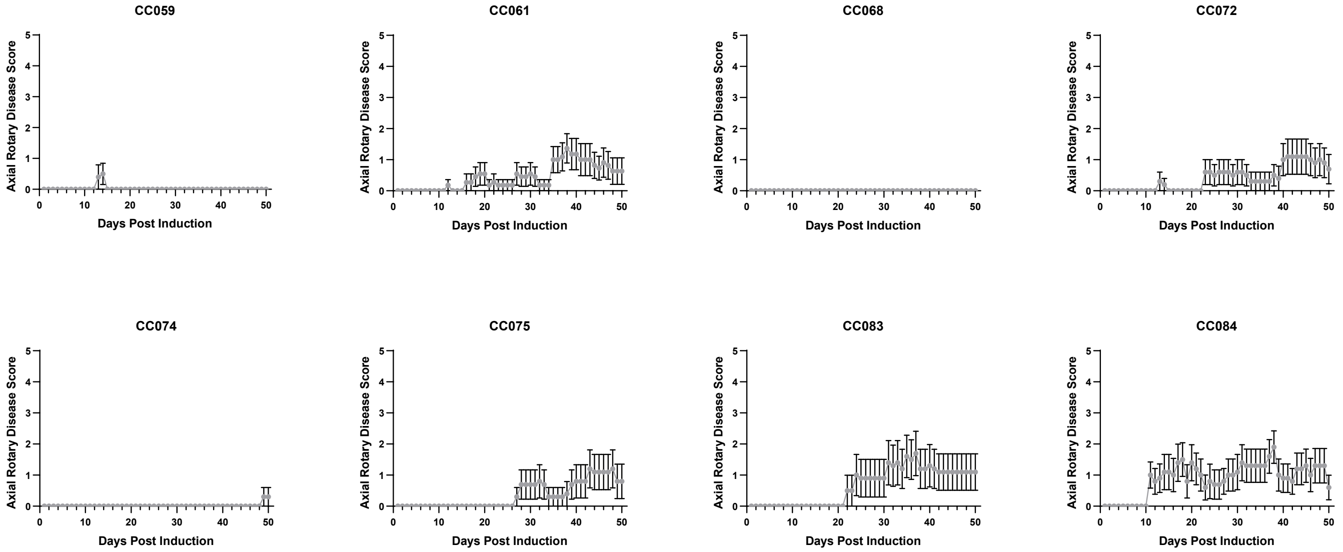


**Figure S6. Axial Rotary EAE disease course profiles for CC strains.** EAE was induced and observed for 50 days in CC mice as described in Figure 1. Post observation, disease course profiles for each strain were derived from daily AR-EAE disease scores (see Materials and Methods). AR-EAE disease course profiles as calculated by strain average for each CC strain are displayed. CC strains are displayed in numerical order by strain number and strains highlighted in Figure 2 retained strain specific label coloring.

### Figure S7 1/2

#### Axial Rotary Disease Course – Sexes

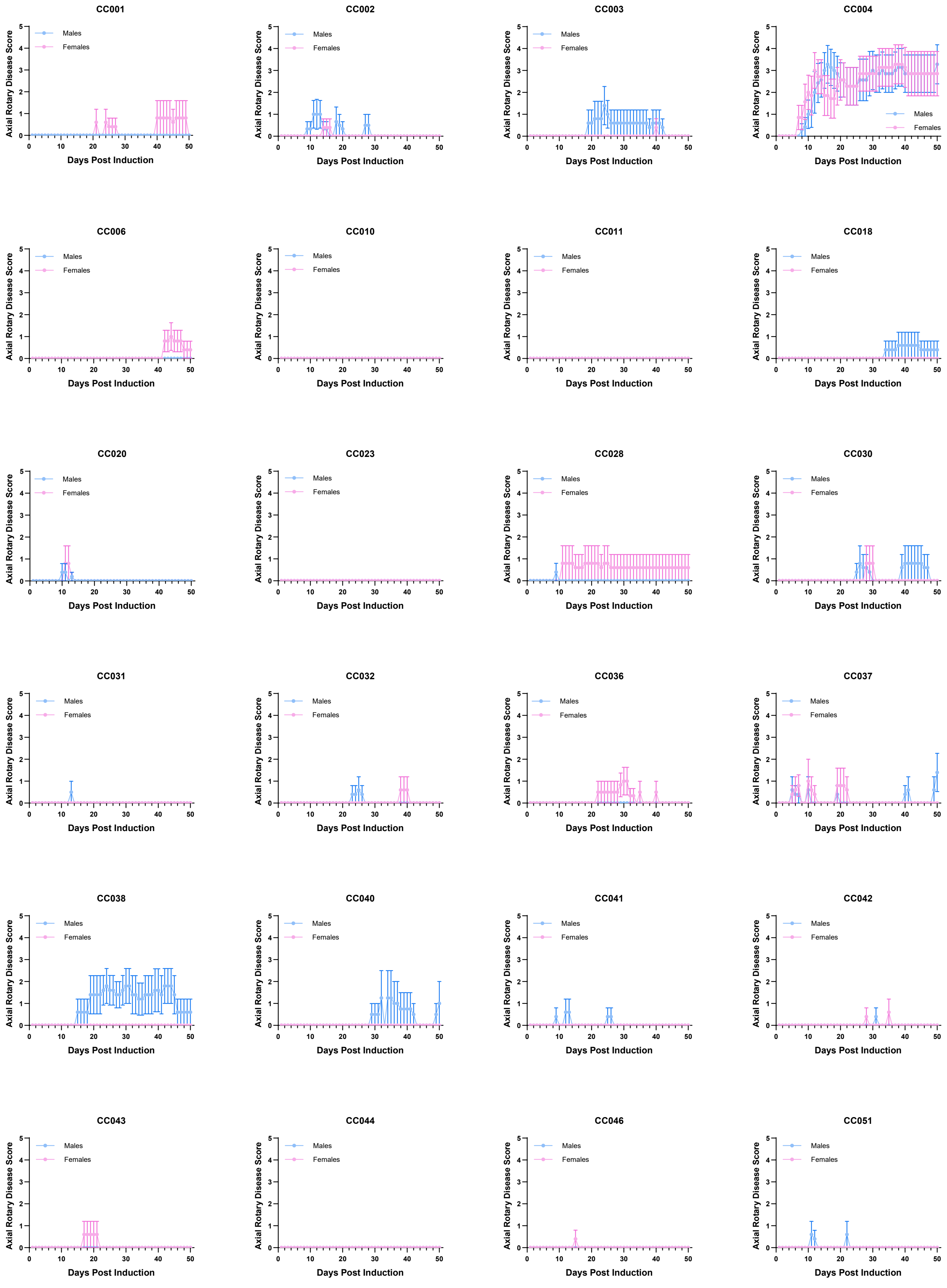

### Figure S7 2/2

#### Axial Rotary Disease Course – Sexes

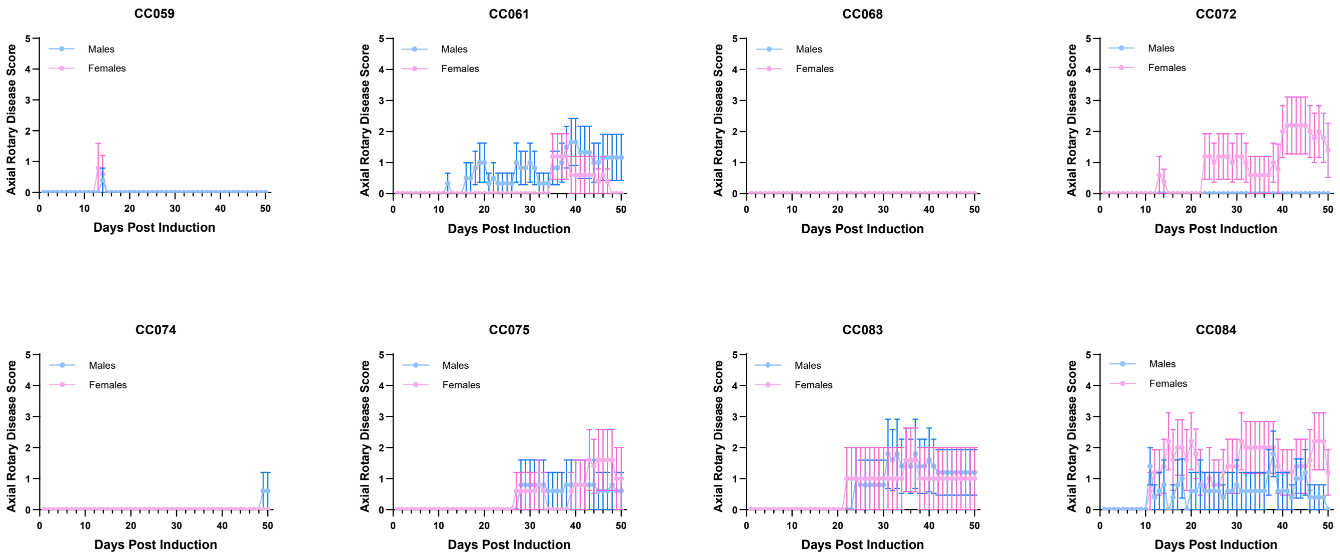


**Figure S7. Axial Rotary EAE disease course profiles for CC strains separated by sex.** EAE was induced and observed for 50 days in CC mice as described in Figure 1. Post observation, disease course profiles for each strain were derived from daily AR-EAE disease scores (see Materials and Methods). Sex specific AR-EAE disease course profiles as calculated by male (blue) and female (pink) averages for each CC strain are displayed. CC strains are displayed in numerical order by strain number.

### Figure S8

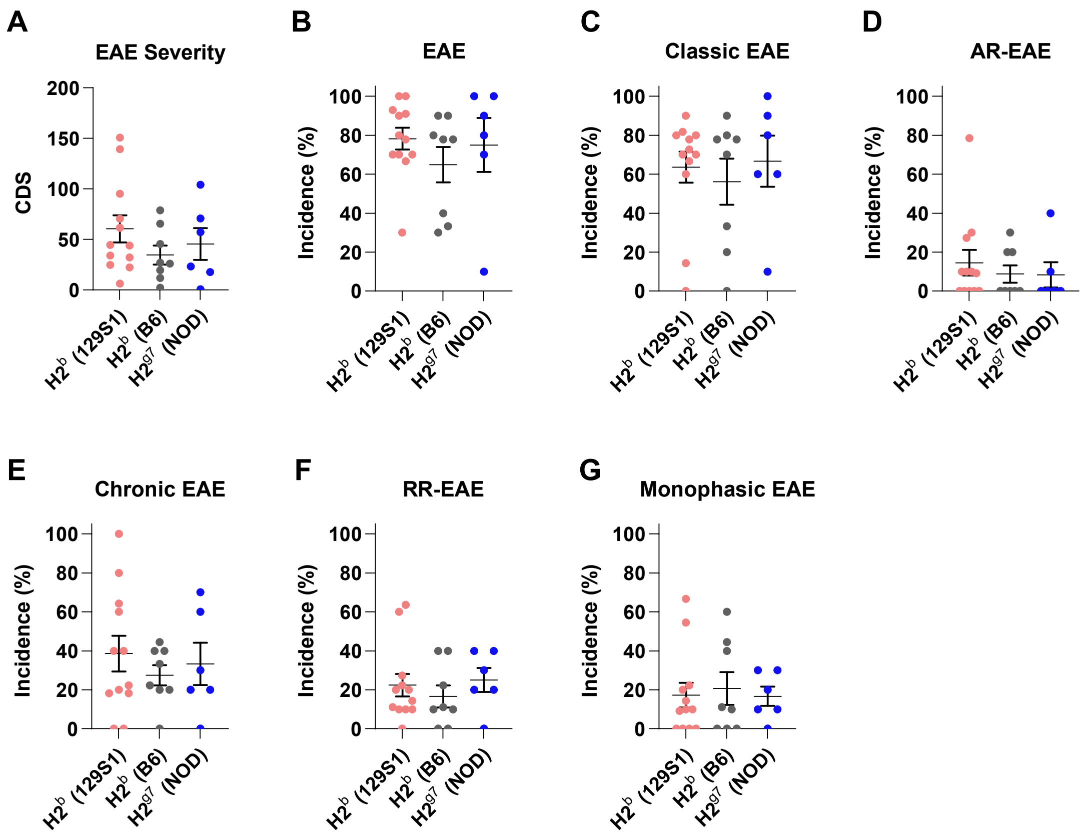


**Figure S8. CC EAE disease phenotypes are independent of CC founder specific H2 effects.** EAE was induced and observed for 50 days in CC mice as described in Figure 1. Mice were observed daily for a total of 50 days starting at 5 days post induction for the presence of clinical disease symptoms which were quantified to assess the overall EAE disease profile as described in Materials and Methods. (**A-G**) Distribution of strain (**A**) cumulative disease score (CDS), (**B**) total EAE incidence, and incidence of (**C**) classic EAE, (**D**) AR-EAE, (**E**) chronic EAE, (**F**) RR-EAE, and (**G**) monophasic EAE, grouped by founder derived *H2*$^{b}$ and *H2*$^{g7}$ homozygous haplotypes. Significance of differences between haplotypes was determined by one-way ANOVA with Tukey's multiple comparisons test and indicated by asterisks where significant.

### Figure S9

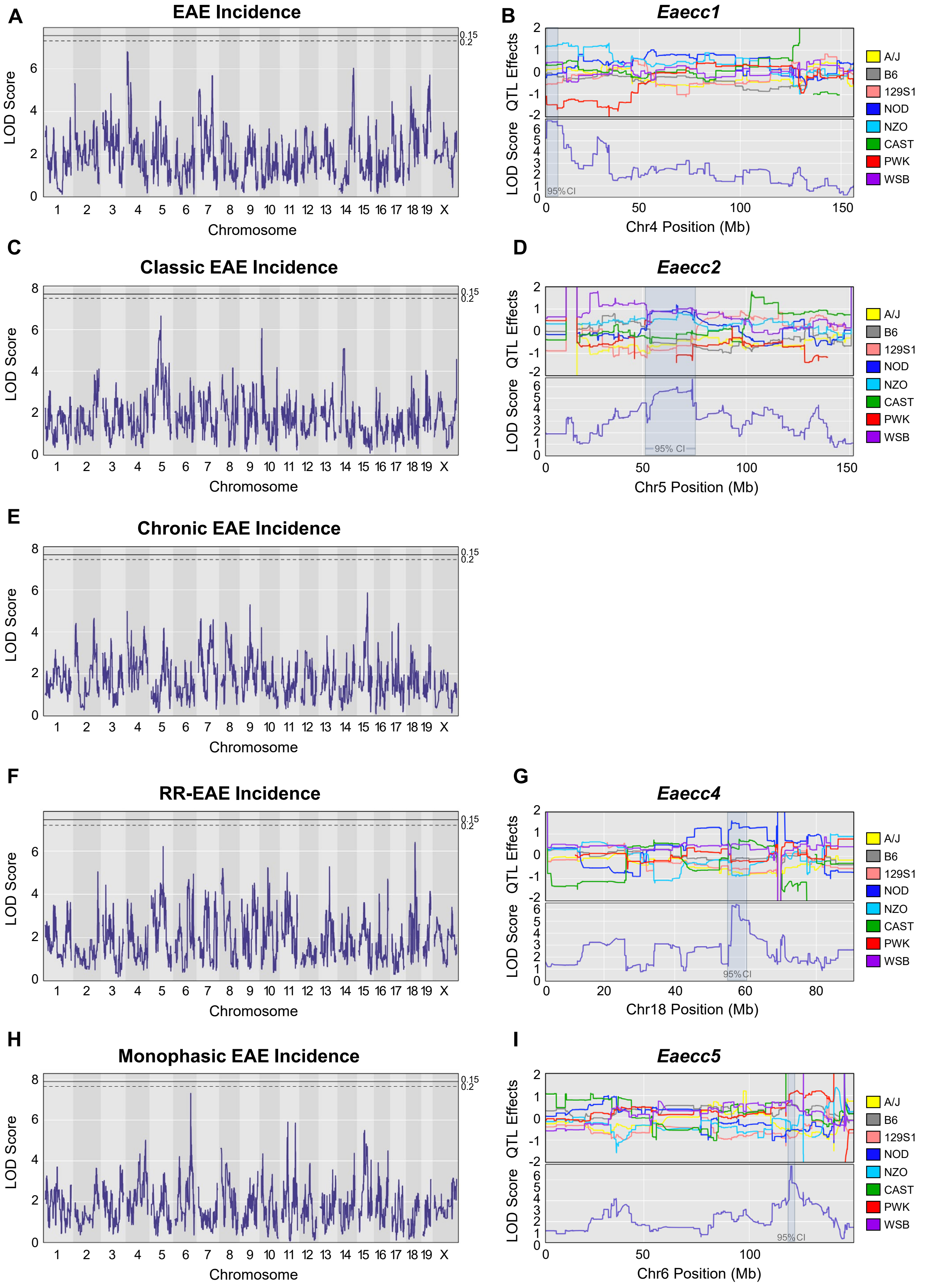

A
EAE Incidence
B
Eaecc1
C
Classic EAE Incidence
D
Eaecc2
E
Chronic EAE Incidence
F
RR-EAE Incidence
G
Eaecc4
H
Monophasic EAE Incidence
I
Eaecc5
LOD Score
Chromosome
QTL Effects
95% CI
Chr4 Position (Mb)
Chr5 Position (Mb)
Chr18 Position (Mb)
Chr6 Position (Mb)
0.15
0.2
A/J
B6
129S1
NOD
NZO
CAST
PWK
WSB

**Figure S9. QTL analysis reveals distinct genetic linkage patterns for several EAE incidence traits.** EAE was induced and evaluated in 32 CC strains, as described in Figure 1. EAE disease phenotypes and quantitative trait variables were calculated, and quantitative trait loci (QTL) mapping was performed (see Materials and Methods). (**A**) Manhattan plot demonstrating logarithm of odds (LOD) traces for QTL mapping assessing genome association with total EAE incidence, and (**B**) corresponding CC founder allele effects plot for lead QTL identified on chromosome 4 -*Eaecc1*. (**C**) Manhattan plot demonstrating LOD traces for QTL mapping assessing genome association with classic EAE incidence, and (**D**) corresponding CC founder allele effects plot for lead QTL identified on chromosome 5 – *Eaecc2*. (**E**) Manhattan plot demonstrating LOD traces for QTL mapping assessing genome association with chronic EAE incidence. (**F**) Manhattan plot demonstrating LOD traces for QTL mapping assessing genome association with RR-EAE incidence, and (**G**) corresponding CC founder allele effects plot for lead QTL identified on chromosome 18 – *Eaecc4*. (**H**) Manhattan plot demonstrating LOD traces for QTL mapping assessing genome association with monophasic EAE incidence, and (**I**) corresponding CC founder allele effects plot for lead QTL identified on chromosome 6 – *Eaecc5*. For Panels (**A**), (**C**), (**E**), (**F**) and (**H**), genome wide significance thresholds of 15% (solid line) and 20% (dashed line) were determined by permutations (n=1000). Panels (**B**), (**D**), (**G**), and (**I**) use the conventional color designations for CC founder strains as follows: yellow: A/J, grey: C57BL/6J, pink: 129S1/SvImJ, dark blue: NOD/ShiLtJ, light blue: NZO/HILtJ, green: CAST/EiJ, red: PWK/PhJ, and purple: WSB/EiJ.

### Figure S10

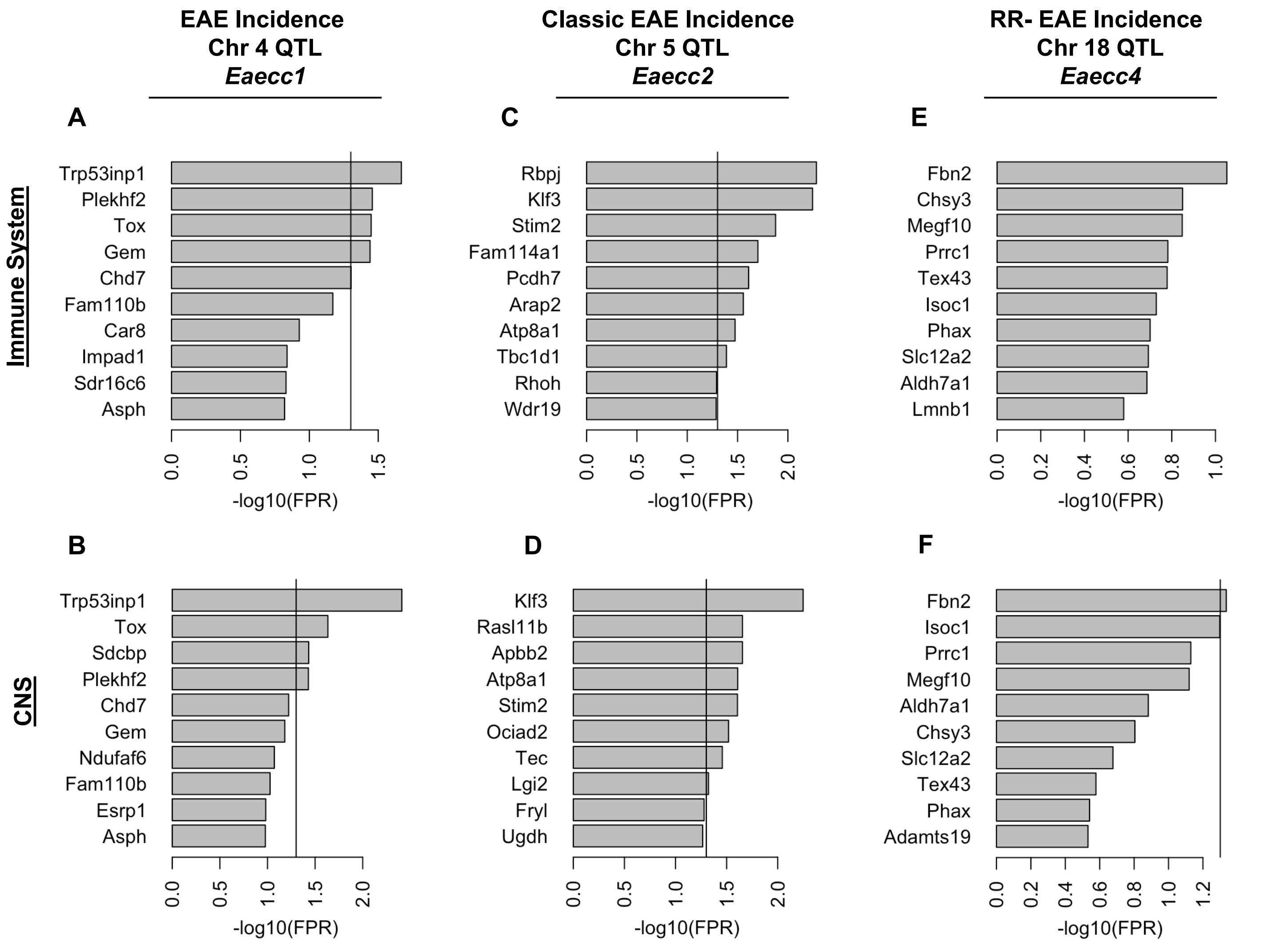


**Figure S10. Functional candidate gene prioritization nominates distinct genes associated with unique EAE incidence traits.** Support vector machine classifiers ranked gene candidates associated with EAE incidence QTLs in the context of either the CNS or immune system as described in Figure 8. Ranked candidate genes for the total EAE incidence QTL on chromosome 4 – *Eaecc1* for the (**A**) immune system network and (**B**) CNS network, classic EAE incidence QTL on chromosome 5 – *Eaecc2* for the (**C**) immune system network and (**D**) CNS network, and the RR-EAE incidence QTL on chromosome 6 – *Eaecc4* for the (**E**) immune system network and (**F**) CNS network. The solid line in panels **A** – **F** corresponds to the FPR threshold of 0.05.
